# Fitness effects of competition within and between species change across species’ ranges, and reveal limited local adaptation in rainforest *Drosophila*

**DOI:** 10.1101/395624

**Authors:** Eleanor K. O’Brien, Megan Higgie, Christopher T. Jeffs, Ary A. Hoffmann, Jan Hrček, Owen T. Lewis, Jon R. Bridle

**Affiliations:** University of Bristol; James Cook University; University of Oxford; University of Melbourne; Czech Academy of Sciences

## Abstract

Competition within and between species can have large effects on fitness and may therefore drive local adaptation. However, these effects are rarely tested systematically, or considered when predicting species’ responses to environmental change. We used a field transplant experiment to test the effects of intra and interspecific competition on fitness across the ecological niches of two rainforest *Drosophila* species that replace each other along an elevation gradient. For the species with the broader elevational range, we also tested for adaptation to the local abiotic and biotic environment. In both species, intraspecific competition reduced productivity more than interspecific competition at the centre of its elevational range, while interspecific competition had a stronger effect at the range edge, where the competing species is more abundant. Local adaptation was detected in the centre of the range of the more widespread species, but only in the presence of intraspecific competition. This study is the first to demonstrate that fitness effects of inter-specific competition increase at ecological margins, while intra-specific competition has more pervasive effects at range centres. This is a key assumption of “tangled bank” models of community evolution and has important implications for predicting the resilience of ecological networks to global change.

## Introduction

Darwin (1859) used the metaphor of the “entangled bank” to describe the way that interactions within and among species structure an ecological community, due to the narrow range of conditions within which a given species can successfully compete with a neighbouring species. Competitive interactions can have large effects on fitness and are therefore likely to drive adaptive divergence within species (e.g. Stuart et al. 2014, Hargreaves et al. 2019). If we assume a trade-off between resistance to antagonistic interactions such as competition, parasitism or predation, and tolerance of abiotic conditions such as temperature or humidity, then antagonistic biotic interactions should narrow a species’ environmental niche by reducing the range of conditions within which it can persist, causing the evolution of ecological specialisation in communities (Kneitel & Chase 2003; Poisot et al. 2011). Such theory predicts that competition within a species (intraspecific competition) should have the strongest effect on fitness at the centre of the species’ distribution but will become less important towards the range edge. By contrast, at species’ margins, a given level of interspecific competition should have a bigger effect on fitness, leading to species’ turnover along ecological gradients. Interspecific competition should therefore increase in its effects on fitness at or beyond the margins of a species’ range, especially where other closely related species (and likely competitors) increase in frequency.

Spatial variation in the effects of competition on fitness should cause local adaptation if these effects are consistent over time, especially where there is a trade-off between competitive success and resistance to abiotic stress. Studies in angiosperms have found that competition can either increase (e.g. Bischoff et al. 2006; Rice & Knapp 2008) or decrease (e.g. Bischoff et al. 2006) the magnitude of local adaptation. In a meta-analysis of field studies, mostly in plants, Hargreaves et al. (2019) found that local adaptation was neither more prevalent nor stronger in the presence of biotic interactions (including competition), compared with cases where biotic interactions were excluded, despite strong effects of biotic interactions on fitness. However, this effect varied with latitude: in tropical environments, local adaptation was more prevalent when biotic interactions were left intact, however this was not the case in temperate environments.

Strong and pervasive effects of competition on fitness may be more likely in low latitude (tropical) ecosystems, where productivity and biodiversity are generally much higher than in temperate ecosystems, where stronger seasonal fluctuations continually reduce or reset biotic interactions (e.g. Coley & Barone 1996; Schemske et al 2009). At a given latitude, the effects of biotic interactions on fitness should also vary along elevational gradients, given their pervasive effect on community structure, species turnover, and species’ ecology, as well as climatic factors (e.g. Rahbek 1995; Körner 2007; Morris et al. 2015). Consistent with expectations along latitudinal gradients, biotic interactions should be a more important determinant of species’ ecological limits at lower elevation (“warm”) margins, compared with high elevation (“cold”) margins where effects of abiotic factors (e.g. temperature, precipitation) are relatively stronger (e.g. Davis et al. 1998; Pearson & Dawson 2003).

Ecosystems where closely related and ecologically similar species replace each other across predictable environmental gradients provide excellent opportunities to test how competition within and among species affects fitness and drives local adaptation, and how these effects change as species approach their ecological limits. In practice however, it is difficult to disentangle the effects of competition from those of abiotic factors (e.g. temperature, precipitation) on fitness or as drivers of adaptive divergence, because different sources of environmental variation are typically correlated (Godsoe *et al*. 2017). Experiments that manipulate competition and abiotic environmental variation independently are therefore essential for understanding how environmental change mediates the evolution of species interactions.

In the Australian tropical rainforest fruit fly *Drosophila birchii*, transplant experiments have demonstrated that abiotic factors alone cannot explain the species’ field abundance across its climatic range, suggesting an important role for biotic interactions. O’Brien et al. (2017) tested how the fitness of families of *D. birchii* varied when virgin flies were transplanted at a fixed density into field cages at 10 sites along each of two elevation gradients and allowed to mate and produce offspring. They found that fitness (estimated by number of offspring produced) in field cages was highest at the warmest, low elevation sites and declined with increasing elevation. This pattern contrasted with patterns of field abundance, where *D. birchii* was rare at low elevation sites but increased in abundance with elevation up to ∼900m asl. However, given these field cages contained only a single species at low density, the effects of competition and other biotic interactions on fitness were not included. The results from these transplant experiments therefore suggested that although abiotic factors (probably cold limits) can explain the upper limit to *D. birchii*’s elevational range, biotic interactions limit population growth at the warm edge of *D. birchii*’s range, leading to a mismatch between cage productivity and field abundance at low elevations. Despite observing substantial genetic variation in fitness of *D. birchii* in field cages (and clinal divergence in productivity under laboratory conditions), O’Brien et al (2017) did not detect any evidence for local adaptation in these field transplants: *D. birchii* families transplanted to their home site did not have higher fitness when compared with those transplanted from the opposite extreme of the species’ elevational range. However, if competitive interactions are a major determinant of fitness across the elevational range of *D. birchii*, local adaptation may only be revealed in an environment that includes these interactions.

In this study, we use a large-scale field transplant experiment to quantify the effects of ecologically-realistic variation in the intensity of intraspecific and interspecific competition on variation in fitness and life history traits of *D. birchii* and its phylogenetically and ecologically-close relative, *D. bunnanda*. In a novel advance on most previous studies, we conduct these assays of density dependent variation in fitness at sites along one of the elevational gradients used in O’Brien *et al*. (2017), which we already know includes the cold and warm extremes of these species’ distributions. In addition, for *D. birchii*, which is found across a broader range of elevations than *D. bunnanda*, we tested for local adaptation by transplanting flies from different populations across their elevational range. This comparison tested whether local adaptation was revealed by comparing responses of flies from different populations to increasing inter and intraspecific competition at different elevations along the gradient.

We tested the following hypotheses:

1. *The fitness effects of competition is strongest at the warm edge of the species’ ranges:* Competition within and between *D. birchii* and *D. bunnanda* will reduce fitness, but the strength of this effect will vary across the elevational gradient. We predict a stronger effect of competition on fitness at lower elevations, consistent with the expectation that antagonistic biotic interactions are more ecologically important at the warm edges of species’ ranges.
2. *Competition within and between species causes species to replace each other along environmental gradients:* Increasing intraspecific competition will have a greater negative effect on fitness than interspecific competition for a given species within cages transplanted into the elevational range where that species has the highest relative abundance (high elevation for *D. birchii*; low elevation for *D. bunnanda*). By contrast, increasing interspecific competition will have a greater effect than intraspecific competition within cages transplanted to elevations where the competitor species has higher relative abundance.
3. *Selection on competitive ability at different elevations causes local adaptation within species*: *Drosophila birchii* reared from populations close to the transplant location will suffer lower reductions in fitness in response to increasing levels of competition than those reared from populations further away on the elevational gradient. At high elevation (where *D. birchii* is most abundant), intraspecific competition will be a stronger driver of fitness and therefore local adaptation. By contrast, local adaptation will be more strongly associated with interspecific competition at lower elevations, where *D. birchii* is outnumbered by the competitor species (*D. bunnanda*), and intraspecific interactions are relatively rare.

## Methods

### Study species

*Drosophila birchii* and *D. bunnanda* (Diptera: Drosophilidae) are both confined to tropical rainforest habitat, with distributions that overlap at lower latitudes and elevations. They are closely related, similar in size and likely to target similar food and oviposition resources at sites where they co-occur. They do not hybridise in the laboratory (personal observation), and there is no evidence for hybridisation in the wild. *Drosophila birchii* has a broader latitudinal range than *D. bunnanda* (van Heerwaarden et al. 2009), and a broader elevational range within latitudes, with *D. bunnanda* confined to warmer sites at low latitudes and elevations. At the sites sampled for this study, *D. bunnanda* typically outnumbers *D. birchii* ∼3:1 in field traps below 100m elevation, then it declines in abundance at higher elevations. The two species have roughly equal abundance ∼400m asl, and *D. bunnanda* is virtually absent above 500m asl, where abundance of *D. birchii* increases, reaching its maximum around 900m asl (Figure 1; Bridle *et al*. 2009; O’Brien 200*et al*. 2017).

**Figure 1.**
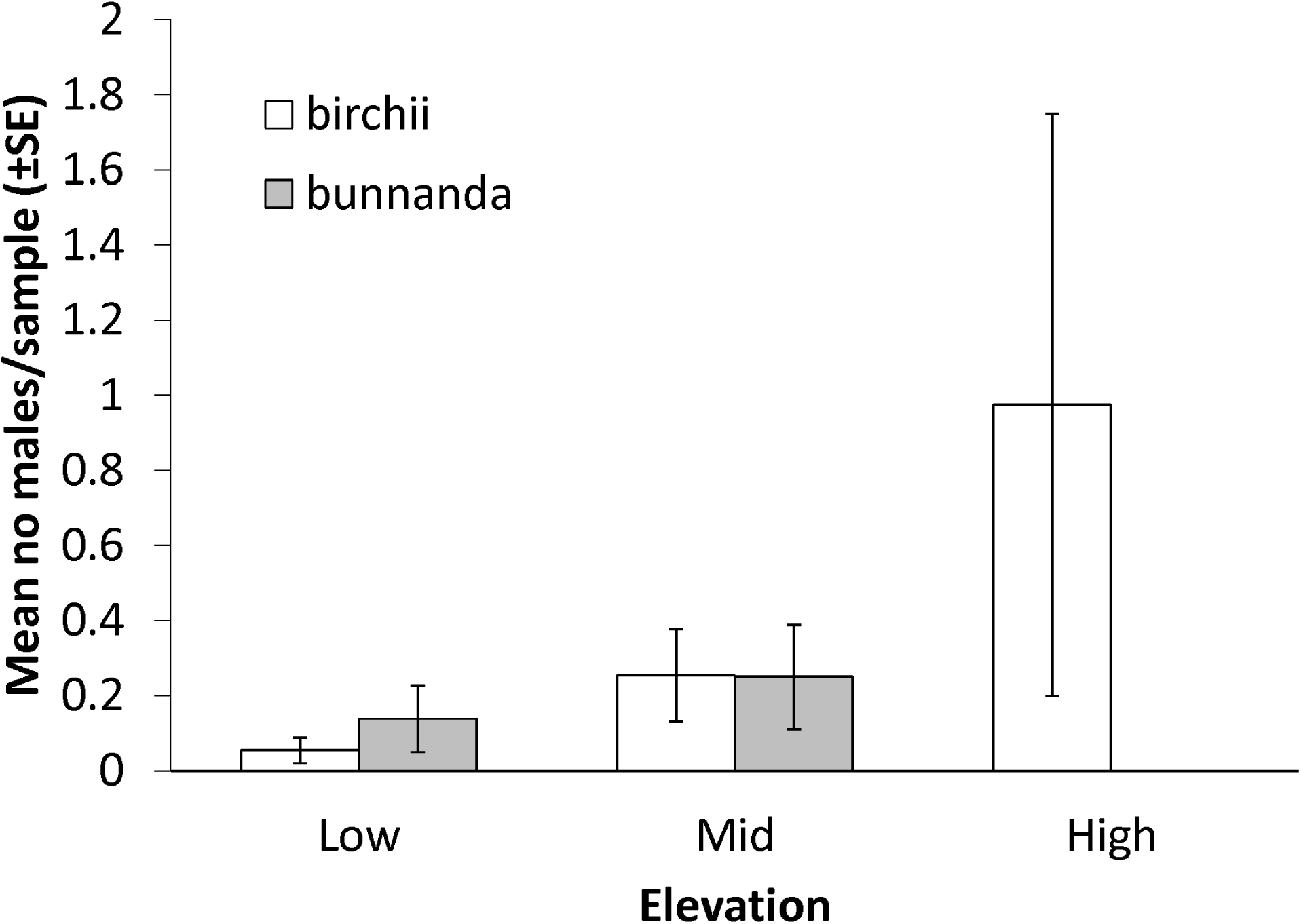
Abundance of *Drosophila birchii* (white bars) and *D. bunnanda* (grey bars) males caught in field traps at Paluma between March – June 2017, including the period when the caged transplant experiment was conducted. Bars indicate the mean number of males of each species caught per trap per day, with error bars indicating standard errors among days.

### Source of flies used in field transplant experiment

In April 2016, we established 80 *D. birchii* isofemale lines from field-mated females collected at two high elevation (∼900m above sea level (a.s.l.)) and two low elevation (<100m a.s.l.) sites at each of two gradients separated by ∼300 km of latitude: Paluma (19°00’ S, 146°14’ E) and Mt Lewis (16°35’ S, 145°19’ E)(Eight sites in total). Each isofemale line was founded by a single, field-mated female. These ten *D. birchii* isofemale lines per site were maintained in the laboratory for 10 months (∼20 generations). Two generations before establishment of the cage transplant experiment, we mixed lines from the same site together to create eight mass-bred *D*.*birchii* populations (Supplementary Note 1).

We established isofemale lines of the competitor species, *D. bunnanda*, using the same method as for *D. birchii*. However, because *D. bunnanda* is absent above 500m, these lines all came from low elevation sites at each of the two gradients where *D. birchii* was collected. We maintained five *D. bunnanda* lines from each of Paluma and Mt Lewis (10 *D. bunnanda* lines in total) in the laboratory over the same period and under the same conditions as for the *D. birchii* lines. We then combined them all to establish a single mass bred population of *D. bunnanda* (Supplementary Note 1).

All isofemale lines and mass-bred populations were maintained at 23 °C on a 12:12hr light:dark cycle prior to establishment of the field experiment.

### Establishment of field transplant experiment

Two generations after mixing, we separated emergees from mass-bred populations by sex under light CO_2_ anaesthesia within 24 hours of emergence every day over seven days, and held them in single-sex vials (maximum density 10 flies). This ensured they were unmated at the start of the experiment, and that all courtship and mating occurred within field cages. We kept them in single-sex vials for a minimum of 72 hours to recover from the effects of CO_2_ before transplant into field cages. The long collection period was necessary to obtain sufficient numbers of flies but meant that experimental flies varied in age from 3 – 10 days when they were put in field vials. To avoid confounding effects of such variation in age, we mixed emergees from different emergence days within each population prior to establishment of field vials.

We transplanted all populations of *D. birchii* and *D. bunnanda* in vials along one of the field elevation gradients where flies were sourced (Paluma). The two gradients from which the original lines were collected have very similar ranges of abiotic conditions (temperature and humidity), which change in the same way with increasing elevation (O’Brien *et al*. 2017). We established transplant cages at three elevations (‘sites’): Low (80m above sea level (a.s.l.)), Mid (450m a.s.l.) and High (900m a.s.l.). The low and high transplant sites included the sites from which the Paluma isofemale lines were sourced. To account for localised environmental heterogeneity within each elevation, we divided each site into five sub-sites (‘blocks’) of roughly equal size, giving 15 blocks in total. Details on variation in the abiotic environment along the gradient are provided in Supplementary Note 2 and Figure S1.

We transplanted flies in 30 ml plastic vials containing 5 ml of standard *Drosophila* media. Vials were closed with a square of muslin secured with a rubber band, which prevented flies from getting in or out, but allowed free air exchange with the outside, meaning conditions inside the vials tracked external temperature. We placed vials in holders constructed from 600 ml plastic bottles with two 135 × 95 mm windows cut out of the sides, ensuring maximal flow of air around the vial openings. We placed between two and four food vials in each bottle and hung bottles from tree branches at a height 1.3 – 1.8 m above the ground. We suspended a 26cm plastic plate upside down on the twine above each bottle to protect vials from rain and encased each bottle in strong wire mesh (20mm square holes) to prevent damage by vertebrates (particularly birds and small mammals).

We transplanted 19 656 virgin flies (11 808 *D. birchii* and 7 848 *D. bunnanda*) in 972 vials at a range of intraspecific and interspecific densities (see ‘competition treatments’ below) into the three transplant elevations in May 2017. We placed flies in vials less than 24 hours before they were installed in the field and left them in vials at their respective transplant sites for 10 days. Therefore, virtually all courtship, mating and egg-laying happened under field conditions. After 10 days, we removed and discarded any surviving flies, and left vials in the field until emergence began. This was 14 days after the establishment of vials at low elevation, 17 days at mid elevation, and 21 days at high elevation. On the day that the first emergence was observed, all vials at that transplant elevation were removed and taken to the laboratory to enable daily emergence to be recorded accurately. Vials were held in a constant temperature room set to the same mean temperature as the elevation at which they had been transplanted, determined using data from the dataloggers inside cages at that site (Supplementary Note 2; Figure S1), on a 12:12 hr light:dark cycle at 60% relative humidity (RH). At all transplant blocks, virtually all larvae had pupated by the time vials were brought in from the field.

### Competition treatments

We used a response surface design (Inouye 2001), which independently varied the numbers of each species and enabled us to estimate the effects of intraspecific vs interspecific competition. We used 10 treatments, each defined by the number of *D. birchii* and *D. bunnanda* in a vial, with a total density of 6, 12, 24 or 48 flies (Figure 2). In a pilot study, we found that the average productivity of *D. birchii* declined with increasing intraspecific density across this range of densities (Supplementary Note 3; Figure S2), demonstrating that it was an appropriate range for detecting competition effects. We also verified that the size of flies emerging from each treatment lay within the size range of field-caught flies at the same elevation (Supplementary Note 4, Figure 3), implying that our competition treatments are ecologically realistic. We introduced equal numbers of unmated males and females (of each species, for mixed species treatments) into each vial, and transplanted five replicate vials of each Population x Treatment combination to each site (one replicate per block).

**Figure 2.**
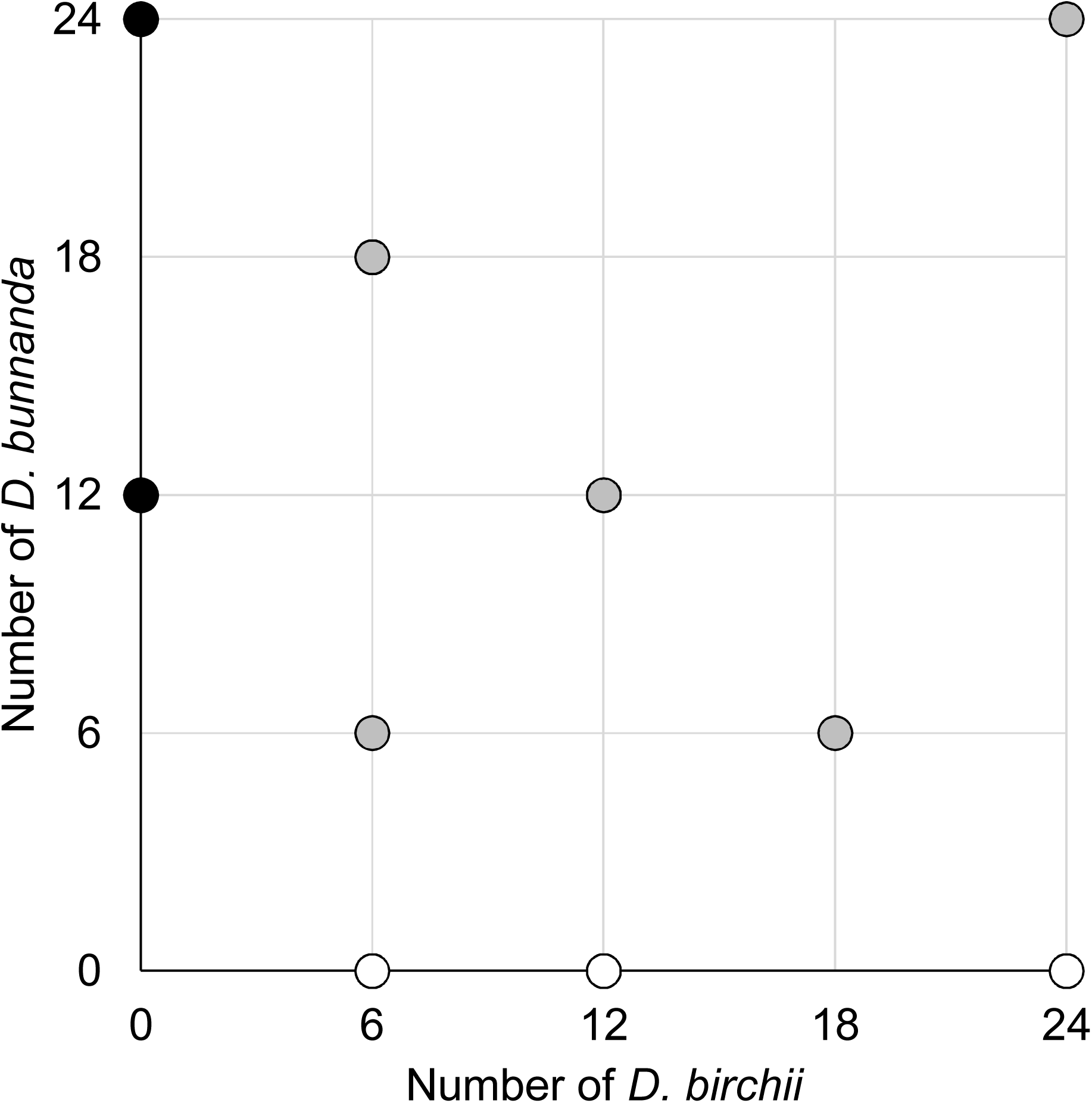
Density treatments in vials, each of which were transplanted to high, middle and low elevation sites at Paluma (average 32 vials/treatment/site). There were 10 treatments, representing different numbers and/or ratios of *D. birchii* and *D. bunnanda*. White circles are treatments with only *D. birchii*, black circles are treatments with only *D. bunnanda*, and grey circles are treatments with a combination of both species.

**Figure 3.**
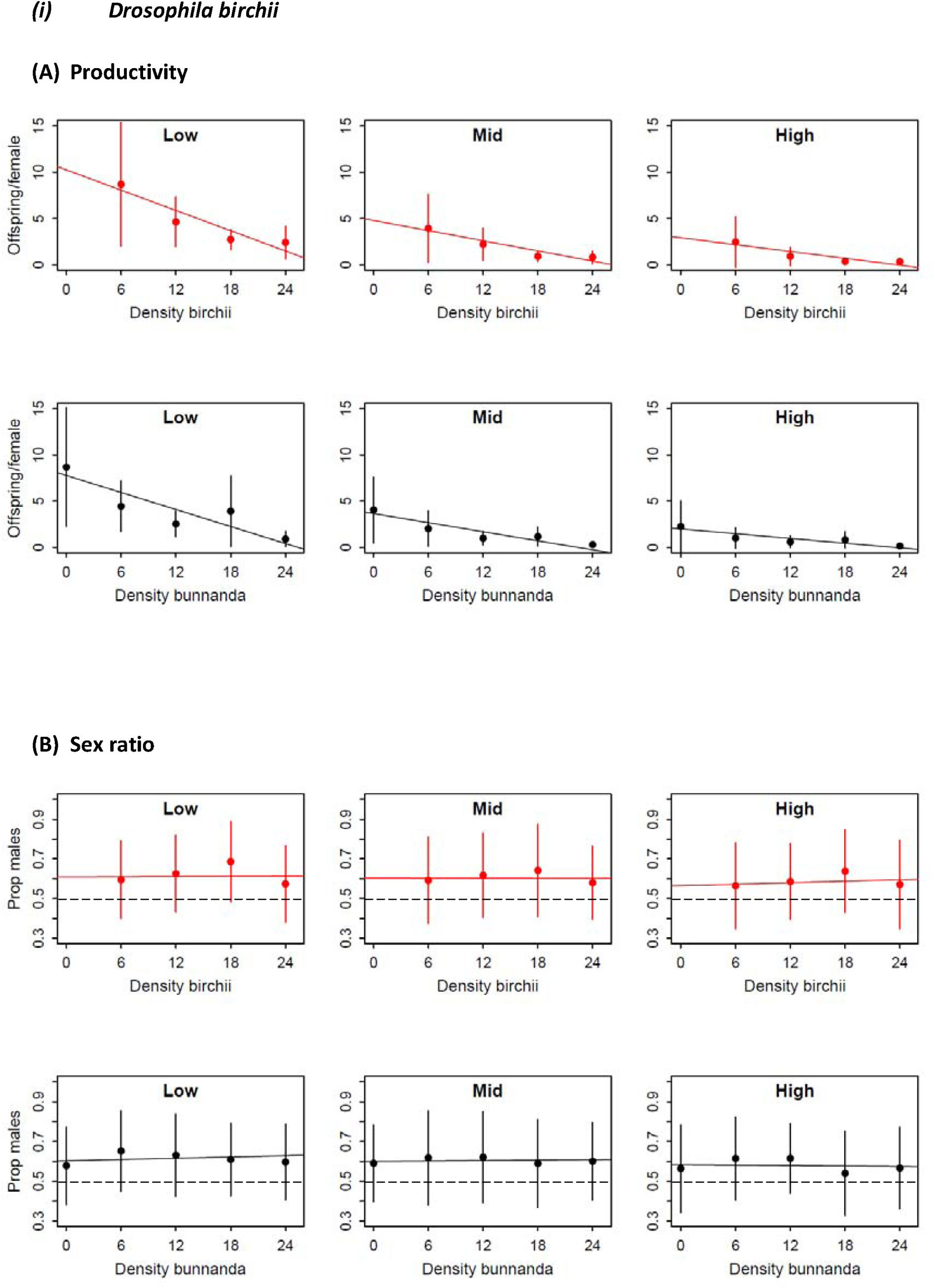
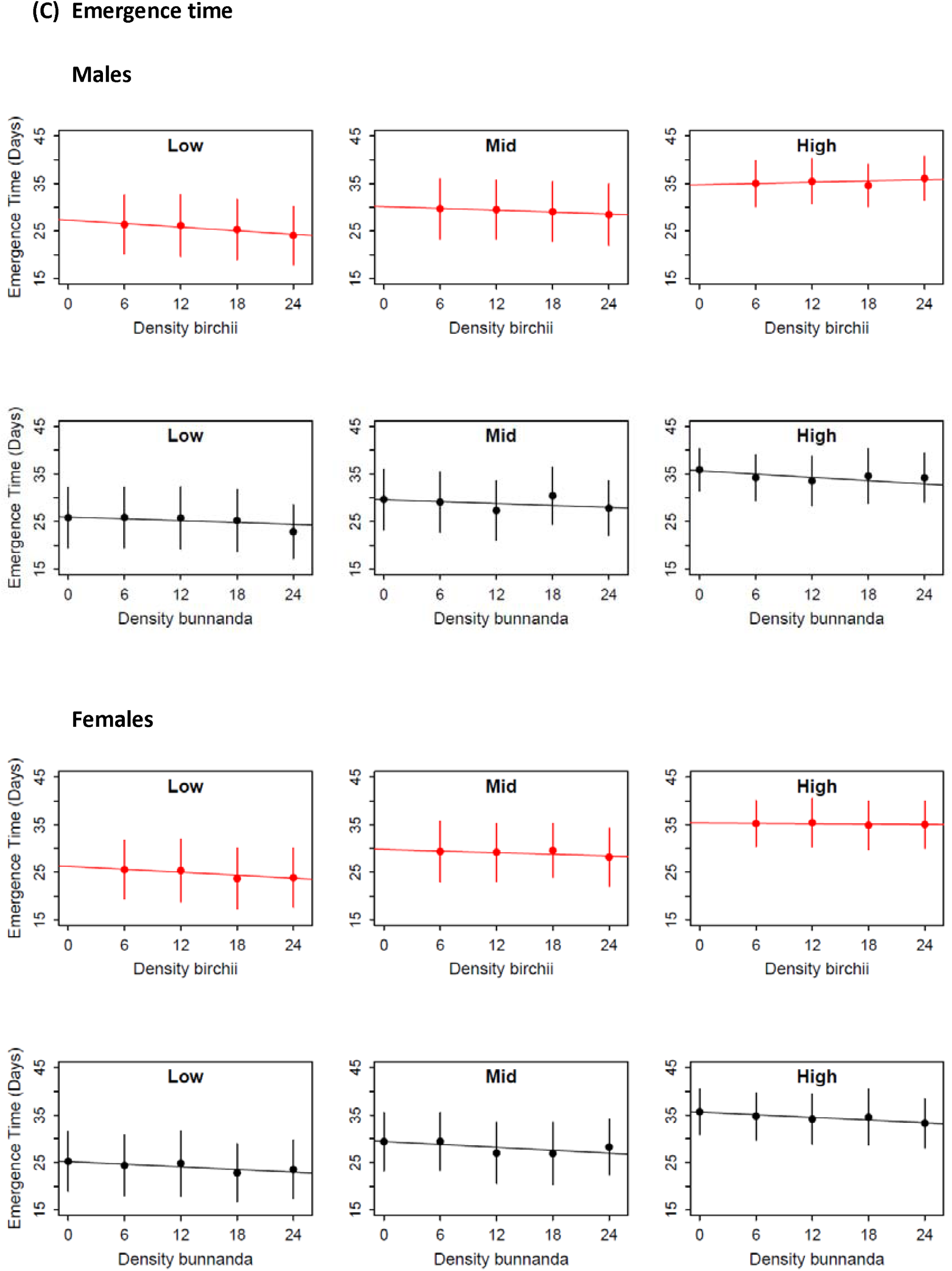
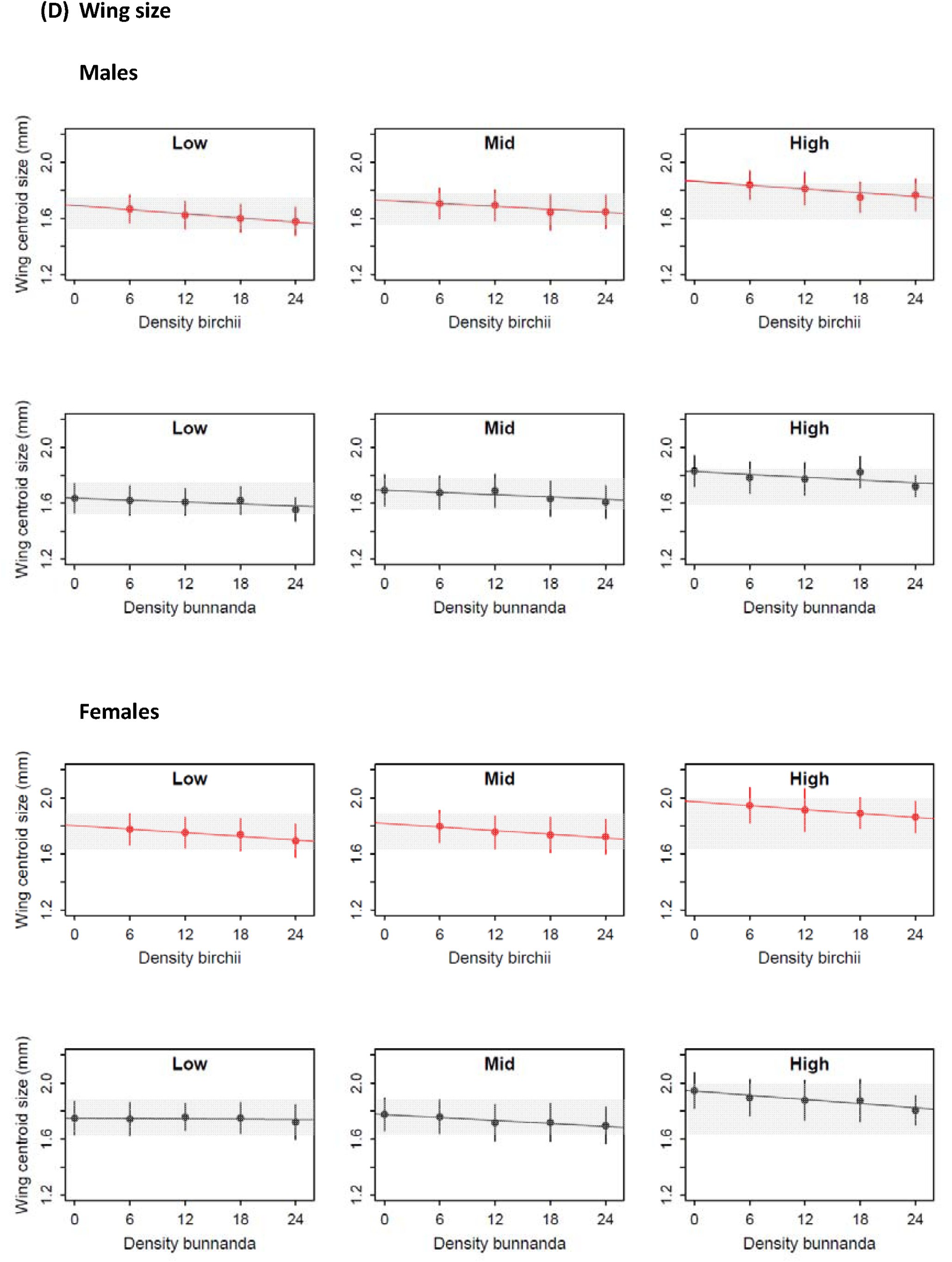
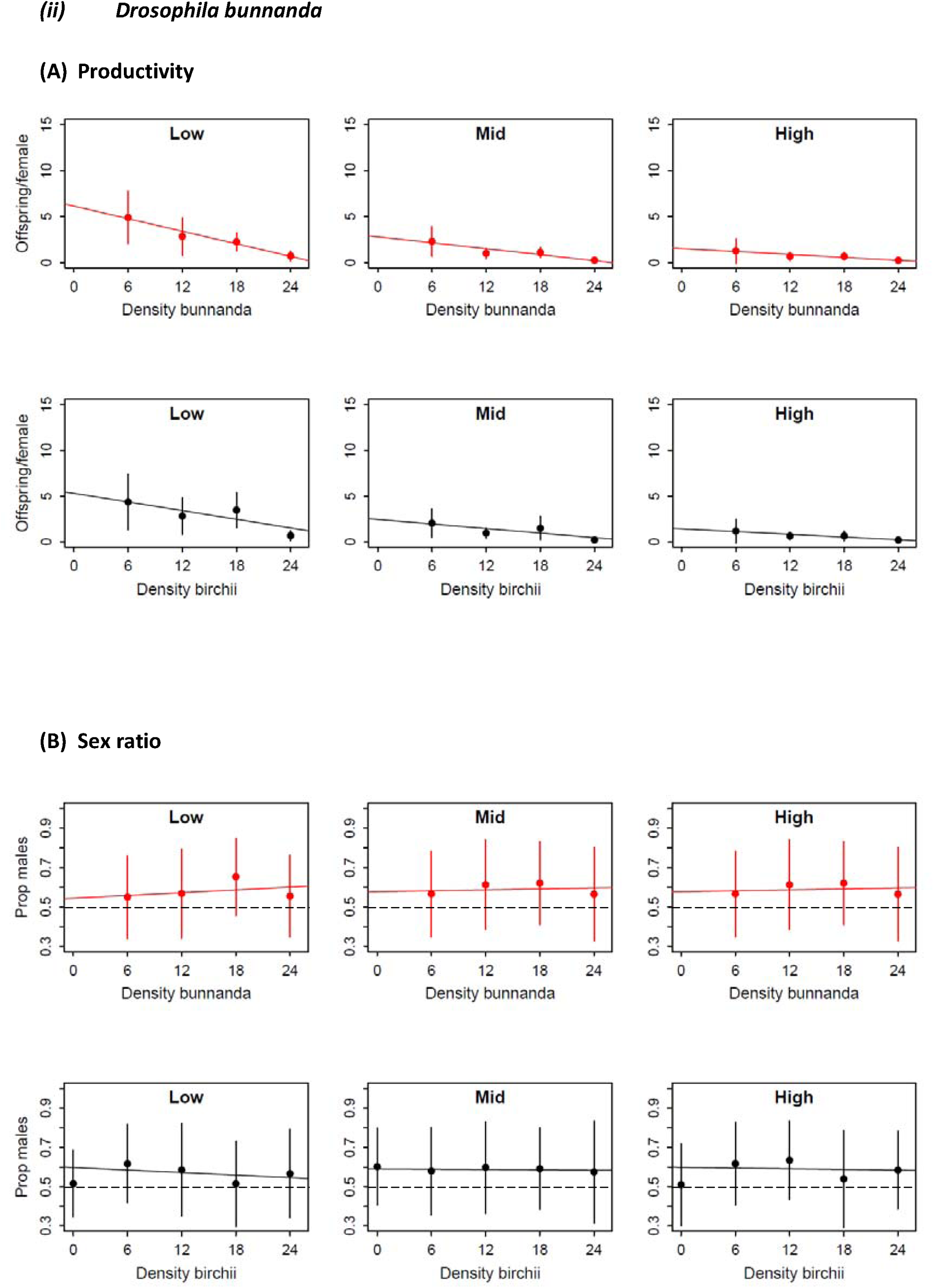
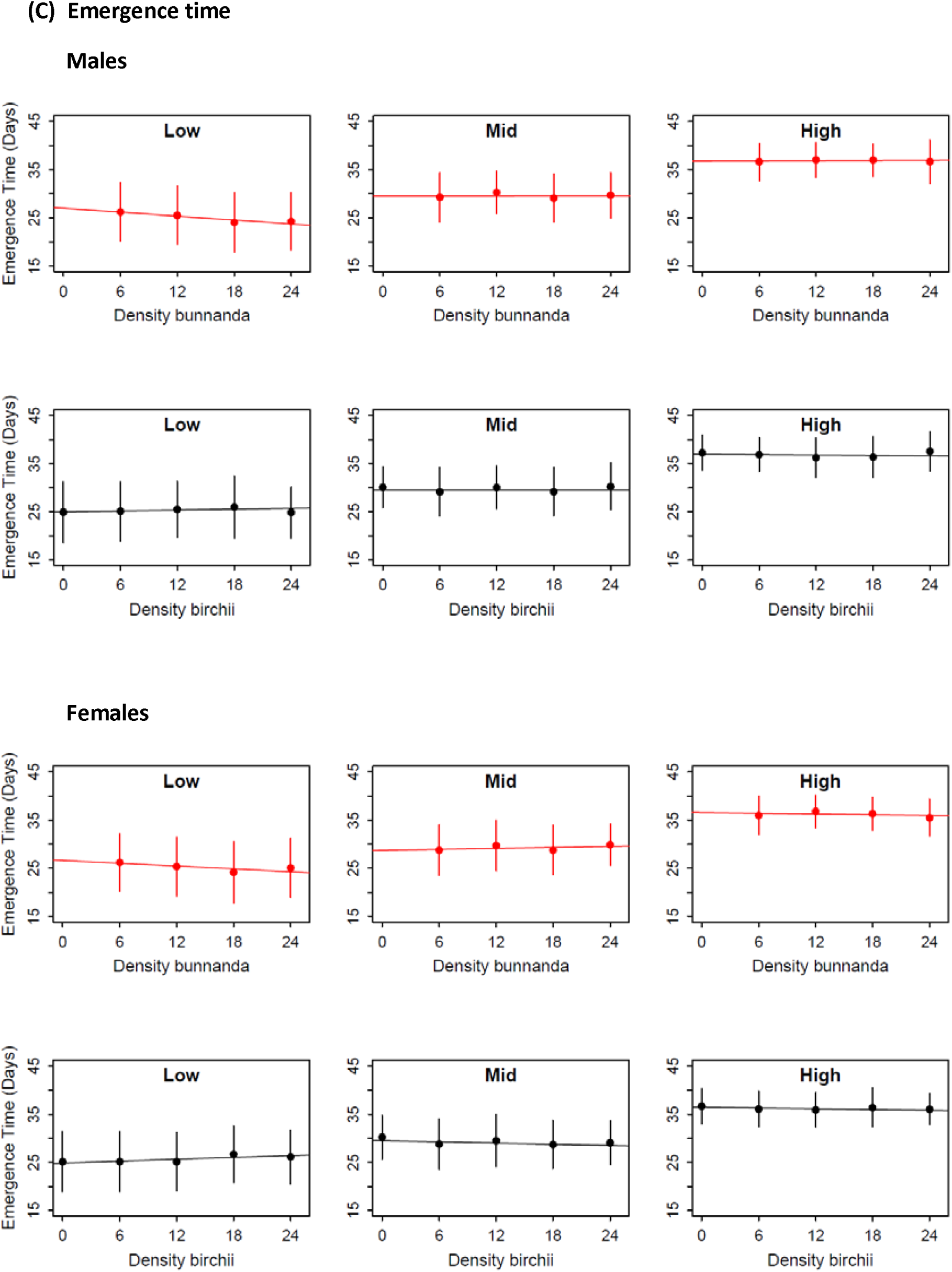
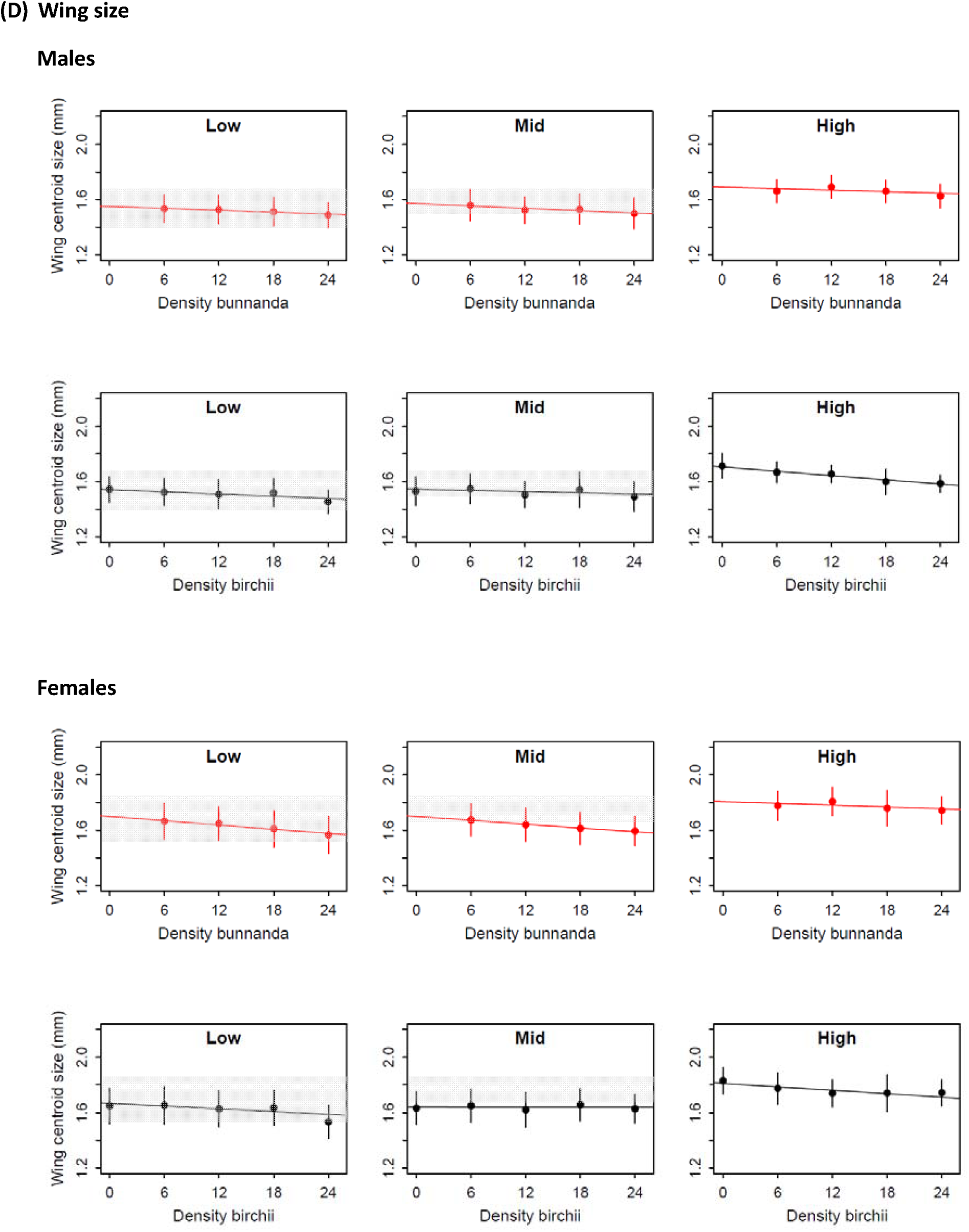
Effect of intraspecific competition (red markers and lines) and interspecific competition (black markers and lines) on productivity, sex ratio, emergence time and wing size of (i) *Drosophila birchii* and (ii) *D. bunnanda* at elevations spanning the abiotic environmental range of both species. Points are means of predicted values from (generalised) linear mixed models that included intraspecific density, interspecific density, elevation and all interactions. Error bars are standard deviations. Lines are the partial regression lines estimated from these models, and therefore represent the independent effect of each type of competition, after accounting for other sources of variation. Where relevant, predicted values have been back-transformed so they are on the original scale. Dashed lines on plots of sex ratio represent an equal sex ratio in the offspring (proportion males = 0.5), therefore values above this line indicate a male-biased sex ratio and values below the line a female-biased sex ratio. The shaded regions on wing size plots show the area bounded by 1 standard deviation either side of the mean wing size of field-caught flies of the relevant sex and species at that elevation.

### Measuring traits of flies emerging from field vials

We removed and counted the number of emergees of each species and sex from each vial on the day emergence began at the transplant site, then daily for the next 10 days and then every three days for an additional nine days to capture any late emergence (20 days total from start of emergence). We undertook species identification and trait measurements blind with respect to treatment or transplant elevation. Male *D. birchii* and *D. bunnanda* were distinguished by their genital bristles (Schiffer & McEvey 2006). Females were identified based on differences in their pigmentation: the dark bands on the dorsal abdomen are straight with sharp edges in *D. bunnanda*, whereas in *D. birchii* they rise in the centre and are more diffuse (M. Schiffer personal communication, and personal observation). For each species emerging from each vial, we recorded the number of male and female offspring emerging each day, to obtain values for the following: (1) productivity (total number of offspring per laying female), (2) offspring sex ratio (number of male offspring as a proportion of the total) for each vial, and (3) emergence time of each fly. We then mounted, photographed and landmarked the right wing of each fly to obtain a measure of (4) wing size, as a proxy for body size. Following the protocol described in Griffiths et al. (2005), we calculated wing centroid size by taking the square root of the sum of the squared distances between each of 10 wing landmarks and the wing centroid.

### Data analysis

#### Testing effects of competition and elevation on traits

We fitted (generalized) linear mixed models to test for the effects of competition (‘Intraspecific/Interspecific density’: see Figure 2 for treatment combinations) and abiotic (‘Elevation’: low, mid or high) environmental variation on each trait. Separate models were fitted for each trait in each species. We applied a Bonferroni correction to adjust the significance level to account for multiple tests. In *D. birchii*, we additionally tested for an effect of the elevation from which the isofemale lines were originally sourced (‘Source elevation’: high or low), and its interactions with competition and transplant elevation. All models were fitted using *lme4* (Bates *et al*. 2015), implemented in *R* v 3.4.2.

For productivity and sex ratio, where vial was the unit of analysis, we fitted models that included fixed effects of intraspecific density, interspecific density, transplant elevation, source elevation and all two-way interactions. We included source population as a random effect. For productivity, we square root transformed data to conform to assumptions of normality, and fitted linear mixed models with the factors described above. For offspring sex ratio, we fitted generalized linear models with a binomial distribution and a logit link function. For emergence time (day) and wing size (mm), which were measured on individual flies, we fitted models with the same fixed and random factors, but additionally included vial as a random effect. Both body size and development time typically differ between males and females in *Drosophila* species (e.g. Santos et al. 1994; Arthur *et al*. 2008), therefore separate models were fitted for each sex. Emergence time and wing size data were normally distributed and were left untransformed.

The significance of fixed and random effect terms in each model was assessed by comparing the log likelihood of a model with or without the relevant term using a chi-squared test. For several traits in both species, the effects of intraspecific and interspecific density varied with elevation (indicated by significant density x elevation terms; Table 1). To further explore the relative effect of intra- and interspecific density on traits within each elevation, we fitted separate linear mixed models for each transplant elevation (i.e. low, mid and high), keeping the remaining fixed and random effects the same as in the full model. For each trait at each elevation, we assessed the relative importance of intraspecific vs interspecific competition in each species using the ratio of their partial regression coefficients, *β*_intraspecific_: *β*_interspecific_ (Anderson & Whiteman 2015).

**Table 1.**
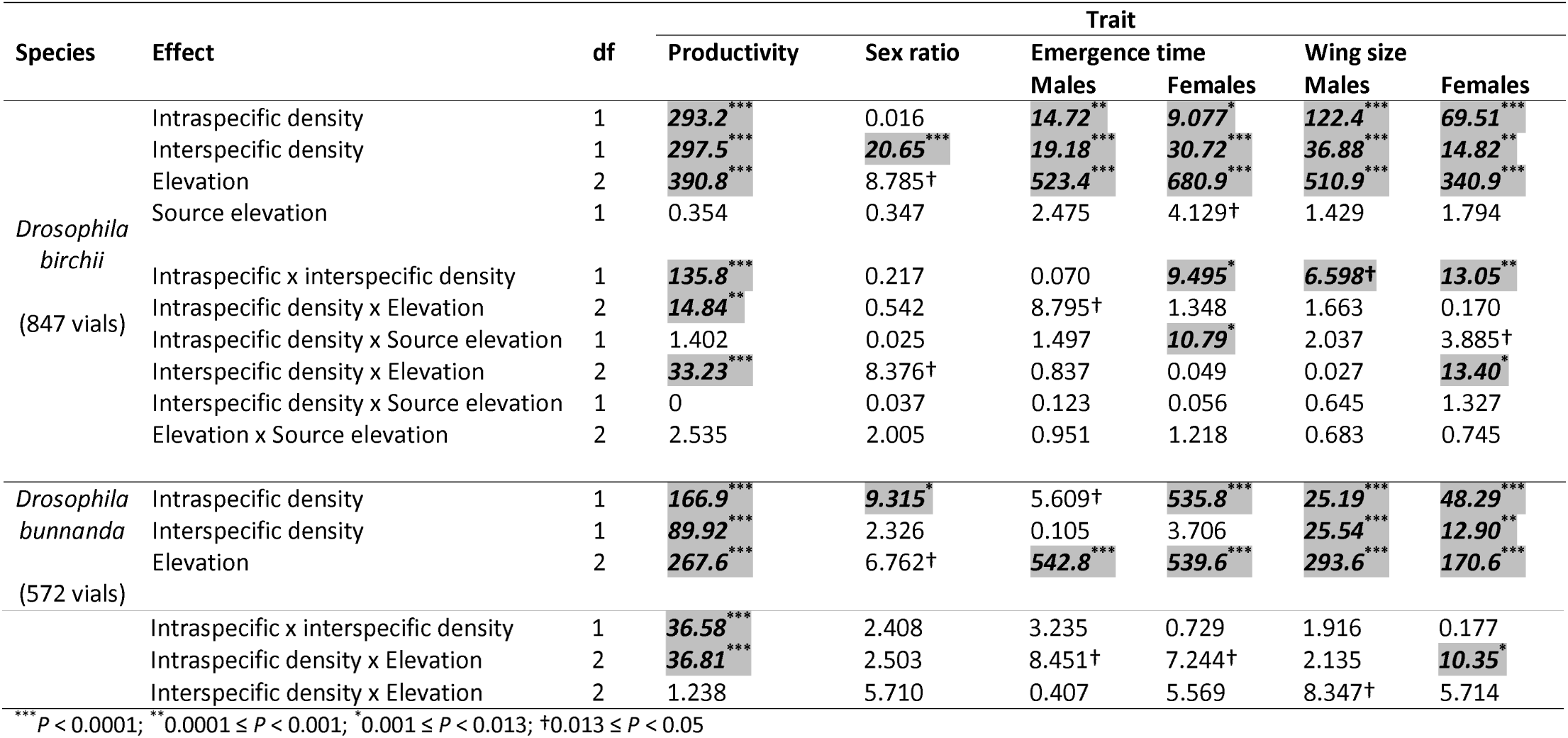
Summary of effects of competition (Intraspecific and Interspecific density) and transplant site (Elevation) on productivity, sex ratio, emergence time and wing size in *Drosophila birchii* and *D. bunnanda*. For *D. birchii*, we additionally tested for divergence between flies sourced from high and low elevation sites (Source elevation). These effects and all 2-way interactions were fitted as fixed factors in (generalised) linear mixed models for each species and trait. For emergence time and wing size, separate models were fitted for each sex. Population was included as a random factor in all models, and vial was additionally included as a random factor in models analysing variation in emergence time and wing size. Values shown for effects on each trait are χ^2^ values obtained from a comparison of the log likelihood of models with and without that fixed effect term included. Within each species, we used a Bonferroni-corrected significance threshold of *P* = 0.013 to test each fixed effect, to account for testing of multiple traits. Tests where the *P*-value for a trait was below this threshold are highlighted and in bold italics. Symbols indicate the range within which *P*-values fall. See legend beneath table.

| Species | Effect | df | Trait |  |  |  |  |  |
| --- | --- | --- | --- | --- | --- | --- | --- | --- |
|  |  |  | Productivity | Sex ratio | Emergence time |  | Wing size |  |
|  |  |  |  |  | Males | Females | Males | Females |
| *Drosophila birchii*<br>(847 vials) | Intraspecific density | 1 | **293.2\*\*\*** | 0.016 | **14.72\*\*** | **9.077\*** | **122.4\*\*\*** | **69.51\*\*\*** |
|  | Interspecific density | 1 | **297.5\*\*\*** | **20.65\*\*\*** | **19.18\*\*\*** | **30.72\*\*\*** | **36.88\*\*\*** | **14.82\*\*** |
|  | Elevation | 2 | **390.8\*\*\*** | 8.785† | **523.4\*\*\*** | **680.9\*\*\*** | **510.9\*\*\*** | **340.9\*\*\*** |
|  | Source elevation | 1 | 0.354 | 0.347 | 2.475 | 4.129† | 1.429 | 1.794 |
|  | Intraspecific x interspecific density | 1 | **135.8\*\*\*** | 0.217 | 0.070 | **9.495\*** | **6.598†** | **13.05\*\*** |
|  | Intraspecific density x Elevation | 2 | **14.84\*\*** | 0.542 | 8.795† | 1.348 | 1.663 | 0.170 |
|  | Intraspecific density x Source elevation | 1 | 1.402 | 0.025 | 1.497 | **10.79\*** | 2.037 | 3.885† |
|  | Interspecific density x Elevation | 2 | **33.23\*\*\*** | 8.376† | 0.837 | 0.049 | 0.027 | **13.40\*** |
|  | Interspecific density x Source elevation | 1 | 0 | 0.037 | 0.123 | 0.056 | 0.645 | 1.327 |
|  | Elevation x Source elevation | 2 | 2.535 | 2.005 | 0.951 | 1.218 | 0.683 | 0.745 |
| *Drosophila bunnanda*<br>(572 vials) | Intraspecific density | 1 | **166.9\*\*\*** | **9.315\*** | 5.609† | **535.8\*\*\*** | **25.19\*\*\*** | **48.29\*\*\*** |
|  | Interspecific density | 1 | **89.92\*\*\*** | 2.326 | 0.105 | 3.706 | **25.54\*\*\*** | **12.90\*\*** |
|  | Elevation | 2 | **267.6\*\*\*** | 6.762† | **542.8\*\*\*** | **539.6\*\*\*** | **293.6\*\*\*** | **170.6\*\*\*** |

|  |  |  |  |  |  |  |  |
| --- | --- | --- | --- | --- | --- | --- | --- |
| Intraspecific x interspecific density | 1 | **36.58**<sup>***</sup> | 2.408 | 3.235 | 0.729 | 1.916 | 0.177 |
| Intraspecific density x Elevation | 2 | **36.81**<sup>***</sup> | 2.503 | 8.451<sup>†</sup> | 7.244<sup>†</sup> | 2.135 | **10.35**<sup>*</sup> |
| Interspecific density x Elevation | 2 | 1.238 | 5.710 | 0.407 | 5.569 | 8.347<sup>†</sup> | 5.714 |

\*\*\* $P < 0.0001$ ; \*\* $0.0001 \leq P < 0.001$ ; \* $0.001 \leq P < 0.013$ ; <sup>†</sup> $0.013 \leq P < 0.05$

#### Testing the power of competition to explain species’ distributions

We tested whether the observed effects of intra and interspecific competition on productivity can explain the relative distributions of *D. birchii* and *D. bunnanda* across the elevation gradient, by comparing predicted productivities with and without competition between the species. We considered three different scenarios: (1) no interspecific competition and constant intraspecific competition across the gradient, (2) no interspecific competition and intraspecific interactions at a frequency proportional to field observations of the abundance of each species, and (3) intraspecific and interspecific interactions at frequencies proportional to the field abundance of each species. For (1), we used the observed productivity of flies in single-species field vials at intermediate density (density = 12). For (2) and (3), we used our field abundance counts (Figure 1) as the starting values for each species’ density (and hence estimates of the frequencies of intraspecific (2 and 3) and interspecific (3) interactions) at each elevation, and used the equations from our elevation-specific models to calculate the expected productivity of each species across the gradient at these densities. Note that because *D. bunnanda* was not found at the high elevation site (density = 0), using this as the starting value meant that the predicted abundance of this species remained zero under each of these scenarios. To enable comparison of observed and predicted abundances under each scenario, we calculated abundances (observed or predicted) of each species at each site relative to the abundance of *D. birchii* at the low elevation site. That is, we set the abundance of *D. birchii* at the low elevation site to 1 and calculated the relative abundance of *D. birchii* at the other sites and *D. bunnanda* at all sites by dividing their observed or predicted abundances by that of *D. birchii* at the low elevation site.

#### Testing for local adaptation

For *D. birchii*, where we transplanted populations from extreme ends of the species’ elevational range, we were able to test for adaptation to the local abiotic or competitive environment. Local adaptation would be revealed by higher fitness of populations in their ‘local’ environment compared with their fitness in other environments (‘home’ vs ‘away’), and/or by higher fitness of local populations than those transplanted from elsewhere in the species’ range (‘local’ vs ‘non-local’)(Kawecki & Ebert 2004). Either of these should result in a source population x transplant environment interaction for fitness. We therefore examined the following interactions in the full models for each trait to test for local adaptation: (i) the ‘Elevation x Source elevation’ interaction to test for local adaptation to the abiotic environment (averaged across all competition treatments); (ii) the ‘Intraspecific density x Source elevation’ and ‘Interspecific density x Source elevation’ interactions to test for adaptation to the local intraspecific and interspecific competitive environments respectively (averaged across transplant elevations and the other form of competition). Furthermore, because abiotic and biotic environmental factors may combine to drive adaptive divergence (if, for example, divergence in competitive ability depends upon the abiotic environment in which it is measured), we examined how Intraspecific/Interspecific density x Source elevation interactions varied across the three transplant elevations. We did not have enough statistical power to detect significant three-way interactions in our full model (i.e. Intraspecific/Interspecific density x Elevation x Source elevation). We therefore used the elevation-specific models and examined: (iii) Intraspecific density x Source elevation and Interspecific density x Source elevation interactions to test for divergence among source populations in their responses to intraspecific and interspecific competition respectively within each transplant elevation. Wherever one of the interactions described above was significant (after Bonferroni correction for multiple comparisons), we examined the pattern of fitness variation to determine whether it was consistent with local adaptation (i.e. superior performance of the ‘home’ or ‘local’ population).

The traits we considered when testing for local adaptation were (i) productivity (since this may be considered a direct measure of fitness) and (ii) wing size, as a proxy for body size, which in *Drosophila* is positively correlated with a range of fitness measures including longevity, female fecundity and male mating success (e.g. Partridge & Farquhar 1983; Santos et al. 1992; McCabe & Partridge 1997).

## Results

### (1) Fitness effects of competition are strongest at the warm edge of the species’ ranges

There was a strong effect of elevation on the overall productivity of both species (Table 1), consistent in direction and magnitude with that observed in *D. birchii* in O’Brien et al. (2017). Mean productivity (averaged across all density treatments) was highest in field cages at the warmest, low elevation site (*D. birchii* offspring per female mean ± SD = 5.52 ± 5.25; *D. bunnanda* mean ± SD = 3.44 ± 2.69; Table S2) and declined with elevation, so that at the high elevation site mean productivity was 24.5% and 27% of that at the low elevation site in *D. birchii* and *D. bunnanda* respectively (*D. birchii* offspring per female mean ± SD = 1.35 ± 2.02; *D. bunnanda* mean ± SD = 0.93 ± 1.05; Table S2).

Emergence time and wing size also varied with transplant elevation (Table 1), with flies of both sexes and species emerging later and at a larger size at higher elevations (Figure 3; Table S2). In *D. birchii*, emergence from cages at the high elevation site was, on average, 9.6 days (males) and 10.4 days (females) later than at the low elevation site (increases of 37.2% and 41.6% respectively; Table S2). In *D. bunnanda*, mean emergence times for males and females respectively increased by 11.7 days (46.4%) and 10.8 days (42.5%) from the low to the high elevation site (Table S2). The mean wing size of *D. birchii* emerging at the high elevation site was 11.0% (males) and 9.7% (females) greater than those emerging at the low elevation site (Table S2). In *D. bunnanda*, these increases were 9.9% and 8.5% for males and females respectively (Table S2). We did not detect an effect of elevation on the sex ratio of offspring of either species (Table 1).

Both intra and interspecific competition greatly reduced the productivity of both species (Table 1; Figure 3) at all transplant elevations (Table 2). The strength of this effect varied with elevation (note the significant interactions of intra and interspecific density x elevation on productivity of *D. birchii*, and intraspecific density x elevation in *D. bunnanda*; Table 1). In both species, the reduction in productivity in response to intraspecific and interspecific competition was greatest at the low elevation transplant site, with weaker effects of both forms of competition at the mid and high elevation sites. In *D. birchii*, the reduction in productivity with each additional competitor of the same species (intraspecific competition; *β*_Intraspecific_) or of the other species (interspecific competition; *β*_Interspecific_) was, respectively, 1.6 and 1.8 times greater at the low elevation site than at the high elevation site. In *D. bunnanda, β*_Intraspecific_ and *β*_Interspecific_ were, respectively, 1.8 and 1.2 times greater at the low than the high elevation site (Table 2; Figure 3).

**Table 2.**
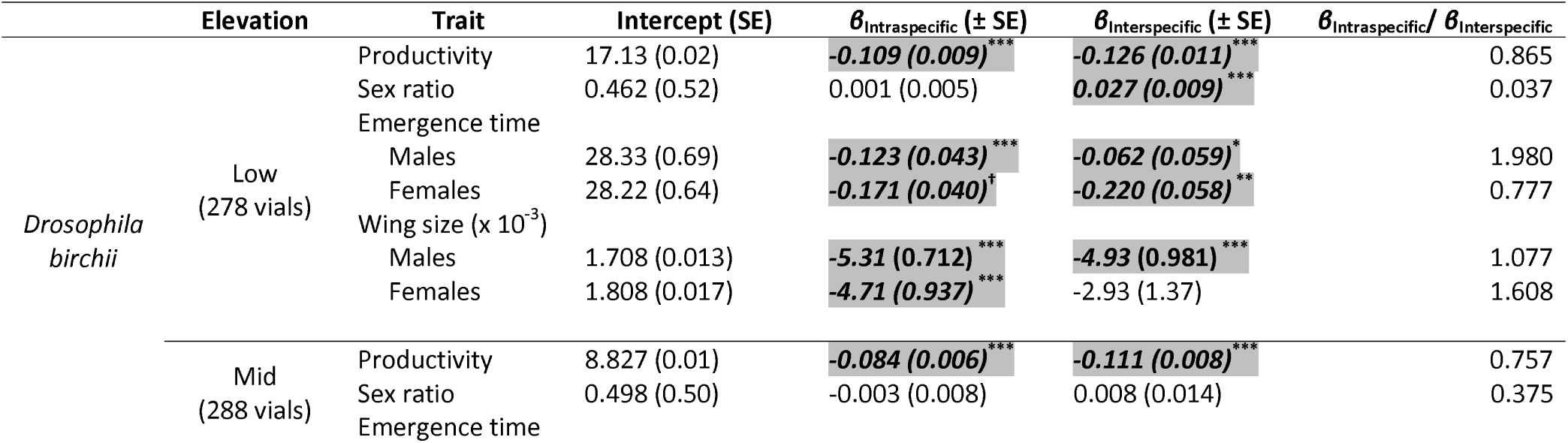
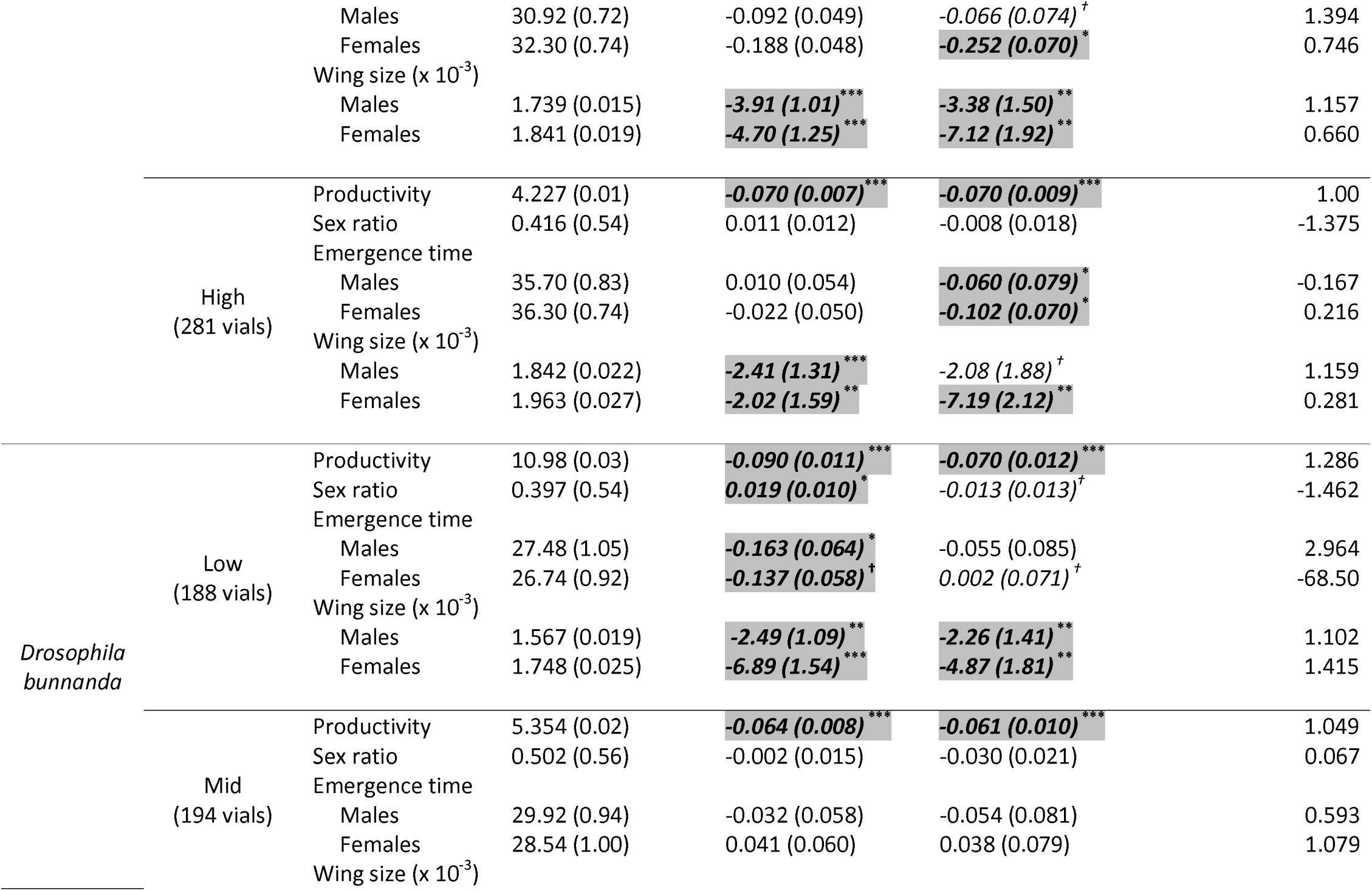
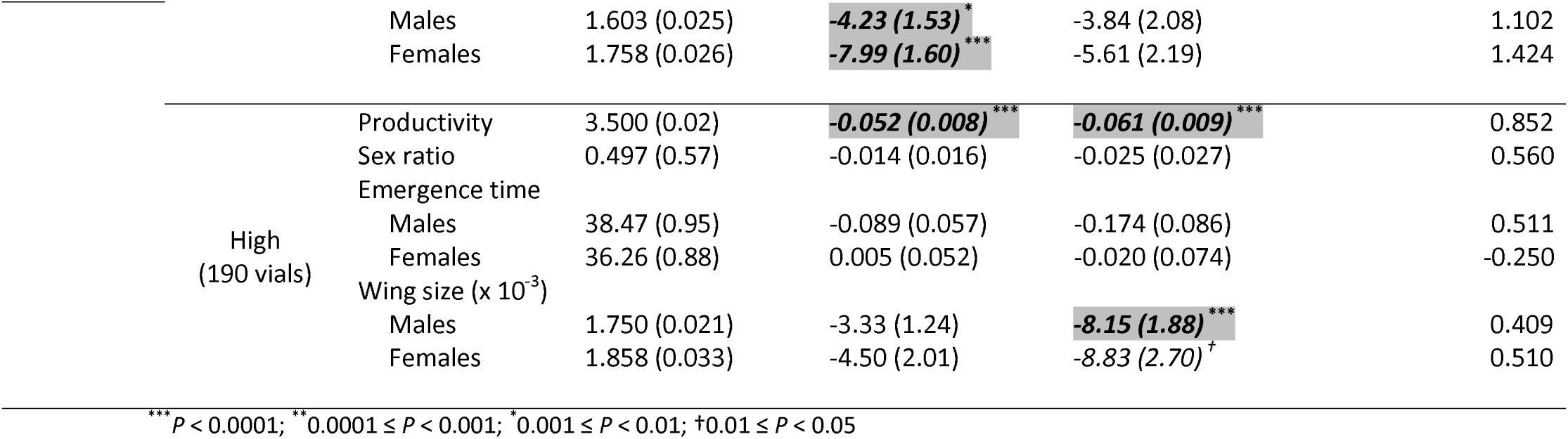
Effects of intraspecific and interspecific competition on productivity, sex ratio, emergence time and wing size in *Drosophila birchii* and *D. bunnanda* at each transplant elevation. Shown are estimates of the partial regression coefficients (*β*) for effects of intraspecific and interspecific density (i.e. holding the other variable constant) on each trait, and its standard error (SE). These were obtained from (generalised) linear mixed models fitted separately for each species, trait and elevation, and separately for males and females for emergence time and wing size. Also shown is the intercept and its SE for each model, back-transformed where necessary so that all values are on the original scale. For productivity and sex ratio, the expected change in the trait value with respect to each predictor variable described by the coefficients (*β*) is on the transformed/link scale used in the model (square root for productivity, log for sex ratio), and so should be interpreted accordingly. The final column gives the ratio of the two partial regression coefficients, which we use to evaluate the relative effect of intraspecific vs interspecific competition on each trait at each elevation. This was calculated as *β*_Intraspecific_/ *β*_Interspecific_, therefore absolute values greater than 1 indicate a stronger effect of intraspecific competition, and absolute values less than 1 a stronger effect of interspecific competition. Negative values indicate that the two types of competition had opposite effects (i.e. one type increased the value of the trait, while the other decreased it. Within each species, we used a Bonferroni-corrected significance threshold of *P* = 0.013 to test each fixed effect, to account for testing of multiple traits. Cases where the partial regression coefficient was significant at this threshold are highlighted and in bold italics. Tests where *P* < 0.05 but not below the corrected threshold are in italics. Symbols indicate the range within which *P*-values fall. See legend beneath table.

There were also strong effects of competition on emergence time and wing size of both species (Table 1). Increasing intensities of intra and interspecific competition resulted in smaller flies that emerged earlier (Figure 3). In contrast to productivity however, the size of the effect of intra and interspecific competition on these traits did not vary with elevation (note lack of significant interactions of intra and interspecific competition with elevation; Table 1). Effects of competition on offspring sex ratio varied between the species and according to the type of competition and were evident only at the low elevation site (Table 2). In *D. birchii*, there was a small but significant increase in the proportion of male offspring emerging as a function of the intensity of interspecific competition at the low elevation site. The mean sex ratio ranged from slightly female-biased (mean prop males ± SD = 0.46 ± 0.02) in field cages where *D. bunnanda* was absent (i.e. no interspecific competition) to slightly male-biased (mean ± SD = 0.54 ± 0.02) in mixed-species cages (Table S2). However, there was no effect of intraspecific competition on sex ratio (Table 1). By contrast, in *D. bunnanda*, we found the opposite: intraspecific competition, but not interspecific competition, increased the proportion of male offspring at the low elevation site (Table 2; Table S2). While the sex ratio of *D. bunnanda* offspring was female-biased in nearly all density treatments at this site (mean prop males ± SD = 0.4 ± 0.18), it ranged from mean ± SD = 0.35 ± 0.07 at the lowest intraspecific density to 0.46 ± 0.02 at the highest intraspecific density (Table S2).

### (2) Competition within and between species causes species to replace each other along environmental gradients

#### Effects of intra and interspecific competition on fitness

In *D. birchii*, interspecific competition had a stronger effect in reducing productivity than intraspecific competition at the low and mid elevation sites (ratios of intraspecific to interspecific density coefficients (*β*_Intraspecific_/*β*_Interspecific_) are less than one; Table 2), where *D. bunnanda* is most abundant (Figure 1). By contrast, at the high elevation site (where *D. birchii* is most abundant and *D. bunnanda* is absent), the negative effects of both forms of competition on productivity were equal (*β*_Intraspecific_:/*β*_Interspecific_ = 1; Table 2).

Similarly, in *D. bunnanda*, intraspecific competition reduced productivity more strongly than interspecific competition at the low and mid elevation sites (*β*_Intraspecific_/*β*_Interspecific_ > 1; Table 2) where intraspecific interactions are most frequent in this species, whereas interspecific competition had a stronger effect in cages at the high elevation transplant site (*β*_Intraspecific_/*β*_Interspecific_ < 1; Table 2) where this species is not normally found.

By contrast, the relative effects of intraspecific vs interspecific competition on other traits of the offspring in the cages (sex ratio, emergence time, wing size) did not show consistent contrasts in effect between the centre and edge of each species’ elevational limits (Table 2).

#### Testing the power of competition to explain species’ distributions

Using the likely frequencies of intraspecific and interspecific interactions at field sites, and the fitness effects of such intensities of interaction within cages to predict species’ relative abundances at each site resulted in predictions that were closer to observed field abundances than predictions made assuming no interspecific competition (Figure S3). Using the productivity of each species in single-species vials to predict field relative abundance gave very misleading results: in particular, it predicted a higher abundance of *D. birchii* (compared to *D. bunnanda*) at the low elevation site and a higher abundance of *D. bunnanda* at the high elevation site (Figure S3B), which is the reverse of what is observed in these ecological communities (Figure 1, Figure S3A). Predictions were significantly improved by assuming intraspecific interactions at a frequency proportional to observed field abundances (Figure S3C), and further by also assuming interspecific interactions occurred at field frequencies (Figure S3D). However, both of these approaches still predicted similar abundances of *D. birchii* at low and high elevations, whereas field observations show much higher abundance of this species at high than at low elevation sites (Figure 1; Figure S3A).

### (3) Selection on competitive ability drives local adaptation within species within a limited part of the elevational range

Evidence for adaptive divergence between high and low elevation source populations of *D. birchii* was limited. Source elevations did not show overall differences in their trait means (no significant effect of Source elevation; Table 1), nor in their fitness responses to the abiotic or competitive environment (Elevation x Source elevation and Intraspecific/Interspecific density x Source elevation interactions were not significant for productivity or body size; Table 1).

High and low elevation source populations of *D. birchii* did differ in the effect of intraspecific competition on wing size, but only at the high elevation transplant site (see Intraspecific density x Source elevation interactions in Table S1; Figure 4). At this site, the reduction in size of male offspring from low elevation source populations as a function of intraspecific density (*β*_Intraspecific_ ± SE = -0.006 ± 0.002; *P* = 1.95 × 10^−4^) was three times greater than that of offspring from high elevation source populations (*β*_Intraspecific_ ± SE = -0.002 ± 0.001; *P* = 0.114) (Table S1; Figure 4), suggesting local adaptation in competitive ability at high vs low elevation, in terms of its effects on body size, at the site where natural density (and therefore intensity of intraspecific interactions) was highest. In females emerging at this site, the difference was even more striking: Intraspecific density reduced mean wing size of female *D. birchii* from low elevation source populations (*β*_Intraspecific_ ± SE = -0.009 ± 0.002; *P* = 5.52 × 10^−6^) by nine times as much as it did wing size of high elevation source populations (*β*_Intraspecific_ ± SE = -0.001 ± 0.002; *P* = 0.439) (Table S1; Figure 4). Source elevations did not differ in their fitness responses to interspecific competition at any of the transplant elevations (Table S1).

**Figure 4.**
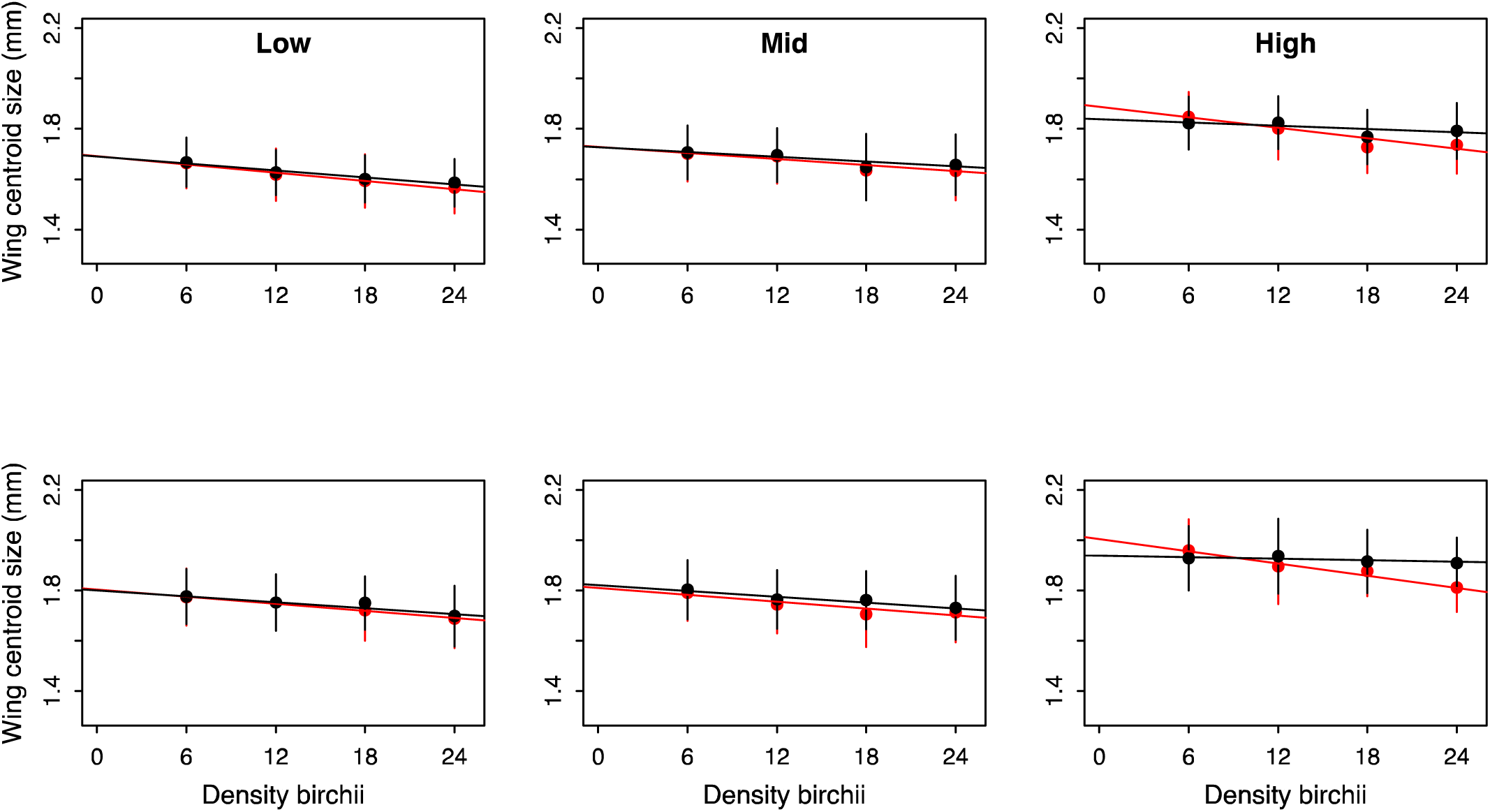
Effect of intraspecific competition (given by the density of *D. birchii*) on wing size of *Drosophila birchii* males (top) and females (bottom) from low elevation (red points) and high elevation (black points) source populations, when transplanted to the low, mid and high elevation sites. Points are means of predicted values from linear mixed models that included intraspecific density, interspecific density, source elevation and all interactions, fitted separately for males and females. Error bars are standard deviations. Lines are the partial regression lines estimated from these models, and therefore represent the independent effect of intraspecific competition, after accounting for other sources of variation. Intraspecific competition caused a significant reduction in wing size of both males and females at all transplant sites. However, the response of low and high elevation source populations only differed at the high elevation site, where intraspecific competition caused a greater reduction in the wing size of flies sourced from low elevation than those from high elevation in both males and females.

## Discussion

### Competition reduces fitness more at the warm margin of species’ ranges

Biotic interactions are thought to be a more important determinant of fitness, and therefore species’ range limits, at lower latitudes (e.g. Coley & Barone 1996; Schemske et al 2009) and elevations (e.g. Davis et al. 1998; Pearson & Dawson 2003). However, tests of this hypothesis have yielded inconsistent results (Moles & Ollerton 2016; Grant et al. 2018). Our results suggest that competitive interactions limit population growth more at the low elevation (warm) edge of the ranges of *D. birchii* and *D. bunnanda*, than at their high elevational limits. The low elevation site also had the highest mean productivity overall. Therefore, for a given interaction density (here determined by varying the number of flies introduced to a cage), larval competition was greatest at the low elevation site, meaning food resources would be depleted more rapidly.

The large reductions in productivity as intra and interspecific competition is increased in cages transplanted to the low elevation site will be compounded by the increased proportions of male offspring at these sites, which will reduce future population growth rate, and therefore evolutionary potential at this range margin (Bridle et al. 2019). Although the effects on sex ratio were subtle, the most prevalent competitive interactions at this site (interspecific for *D. birchii*; intraspecific for *D. bunnanda*) both increased the proportion of offspring that were male. Males are smaller than females in both of these species (Figure 3), and presumably less costly to produce. Therefore, skewing the sex ratio towards males when faced with competitive stress may be a strategy for maximising the number and fitness of offspring produced, consistent with optimal sex allocation theory (Trivers & Willard 1973). It has been shown that female *Drosophila melanogaster* can adjust the sex ratio of their offspring in response to the age of their mate (Mange 1970; Long & Pischedda 2005), and that this may be adaptive. It is not known whether *D. birchii* and *D. bunnanda* are also able to actively manipulate the sex ratio of their offspring, or whether the effect on sex ratio we observe is due to higher survival of male offspring during development. This will be explored further in a future study.

### *Competition shapes the relative distributions of* D. birchii *and* D. bunnanda

The extent to which competitive interactions shape species’ distributions, and under what conditions, is a longstanding question in ecology (e.g. Wisz et al. 2013; Godsoe et al. 2015). Our results demonstrate that both the abiotic environment and the intensity of intra and interspecific competition determine the fitness of *Drosophila birchii* and *D. bunnanda* transplanted in cages across their entire elevational ranges. The effects of competition intensity on productivity in cages varied across the elevation gradient in ways that were consistent with the species’ relative distributions, and with the predictions of the “Tangled Bank” theory of community assembly (REF): each species suffers a greater loss of fitness (productivity) due to intraspecific competition within the centre of its range, and due to interspecific competition at its ecological margins, where the competitor species is more abundant in nature.

We used the site-specific intra and interspecific competition effects estimated from our field transplant experiments to predict relative abundance of *D. birchii* and *D. bunnanda* along the gradient to test for evidence that competitive interactions limit the distributions of these species. Including both types of competition resulted in predicted relative abundances that much more closely matched observed abundance in the field, particularly when compared with predictions made assuming constant intraspecific competition and no interspecific competition (Figure S3). This is consistent with conclusions from a previous transplant study in *D. birchii* that the abiotic environment alone cannot explain its elevational distribution, and that biotic interactions are an important limit to population growth, particularly at low elevations (O’Brien et al. 2017). However, our best predictions still over-estimated the relative abundance of *D. birchii* at the low and mid elevation sites, suggesting that additional factors are required to explain the lower range limit of this species, potentially including other competitors, availability of food resources, pathogens and parasitoids. It is for example known that rates of parasitism by parasitoid wasps on *Drosophila* species in these communities increase at lower elevations (Jeffs et al. 2020), and the effect of this on fitness is being assessed in ongoing work.

### Evidence for local adaptation is strongly dependent on the abiotic and competitive environment

Despite the very large fitness effects of the abiotic and competitive environments tested in these transplant experiments, evidence for local adaptation in *D. birchii* along the elevational gradient could only be detected under a limited set of abiotic and biotic conditions. Increasing within-species (intraspecific) competition in cages had a negative effect on productivity and wing size of *D. birchii* at all transplant elevations. However, at the high elevation transplant site, the effect on wing size varied according to the elevation from which isofemale lines had been sourced, with flies from low elevation source populations showing a much greater (3x in males and 9x in females) reduction in mean wing size than high elevation flies in response to increasing intraspecific competition. Given that the natural abundance of *D. birchii* increases with elevation, this result is consistent with local adaptation of high elevation flies to higher intraspecific density (e.g. by the evolution of increased efficiency at extracting nutrient resources in the presence of conspecifics). In female *Drosophila melanogaster*, large body size has been shown to be strongly predictive of fitness (survival and lifetime productivity), but only when tested under cool (not warm) conditions (McCabe & Partridge 1997). If the same is true in *D. birchii*, selection may favour the maintenance of large body size at the cool edge of the range, even in the presence of high intraspecific competition.

In a study of damselflies, Siepielski et al. (2016) also found that source populations differed in their susceptibility to the negative effects of intraspecific density. In contrast to our result, they observed that populations transplanted to their local site showed more, not less, reduction in fitness in response to intraspecific competition. They attribute this to local populations being better adapted to the abiotic environment at the transplant site and therefore more productive, which exacerbated the intensity of competition (Siepielski et al. 2016). However, in our study we did not find any difference in mean productivity of populations from elevational extremes.

Detecting local adaptation in *D. birchii* is therefore strongly dependent on both the abiotic and biotic environment in which it is measured. This likely explains why neither population divergence nor local adaptation was detected in a previous experiment where *D. birchii* sourced from the same elevational range was transplanted at very low density (O’Brien et al. 2017). Our finding that intraspecific competition can strongly affect the likelihood of detecting population divergence (and possible local adaptation) contrasts with that of Hargreaves et al. (2020), who found in a meta-analysis that maintaining biotic interactions in field transplant experiments did not increase the likelihood or strength of local adaptation detected, compared with studies where biotic interactions were excluded. However, it may be that variation among sites in the extent to which biotic interactions reveal local adaptation, such as we observed here, means that such effects are not detectable when averaged over a wide range of environments.

Some of the most compelling evidence for effects of biotic interactions on adaptation comes from experimental evolution studies of microbial communities (e.g. Lawrence et al. 2012; Fiegna et al. 2015; Jousset et al. 2016; Hall et al. 2018; Scheuerl et al. 2020). Such studies have shown that competitive interactions can constrain adaptive responses to the abiotic environment (e.g. Hall et al. 2018) and that individual species evolve at a slower rate when they are maintained in diverse communities than when evolved alone (Fiegna et al. 2015; Scheuerl et al. 2020). This may be because negative biotic interactions such as competition reduce population sizes and therefore evolutionary potential, or due to trade-offs between adaptation to multiple interacting species or between biotic and abiotic adaptation (Barraclough 2015). It is not yet known whether the tendency for species interactions to reduce evolutionary responses generalises to communities of other types of organisms, but if it does, we would expect local adaptation to be weaker at low latitudes and elevations, where species diversity is typically higher (e.g. Rahbek 1995; Hillebrand 2004; Schemske et al 2009). The greater population divergence of *D. birchii* in response to competition at the high elevation site (compared with the low elevation site) appears to support this pattern.

### Conclusions

Using a novel field transplant design, we assessed the fitness effects of competitive interactions between two species of tropical rainforest *Drosophila* (*D. birchii* and *D. bunnanda*) at sites along an elevation gradient spanning the full climatic extent of their distributions. Consistent with expectations from patterns of biodiversity along elevational and latitudinal gradients, we found that fitness effects of both intra and interspecific competition increased towards the warm, low elevation range margin in both species. In each species, intraspecific competition reduced fitness more than interspecific competition at the centre of the species’ distribution, whereas the reverse was true at the margins where the competitor species becomes more abundant, consistent with adaptation to the abiotic environment inhabited by each species. We also detected adaptation to the local competitive environment within *D. birchii*, the more widespread species, but only at the high elevation (cold) end of its distribution, suggesting evolutionary responses are contingent upon both the abiotic and biotic environment. Our findings highlight the importance of considering biotic interactions when investigating limits to species’ distributions and predicting ecological and evolutionary responses to environmental change. This will be particularly important with climate change, which is expected to have profound effects not just on the abiotic environment but on community composition, and therefore the type and frequency of interactions between organisms (Lurgi et al. 2012).

## Acknowledgements

Thank you to Marcus Lee and Giovanni Bianco for assistance with running the field transplant experiment, and to Jenny Cocciardi and Rebecca Moss for maintaining the Drosophila isofemale lines. For assistance with mounting, photographing and landmarking wings, we would like to thank: Katie Andrews, Sasha Anisman, Eliane Belben, Hannah Bray, Guy Burstein, Jack Challice, Amber-Rose Cooper, Lydia Davies, Peter Dobra, Jessie Fernando, Gemma Glasscock, Dunia Gonzales, Vilhelmiina Haavisto, Sophie Harding, Maxwell James, Bertie Loyd, Cameron Matthews, Harry New, Catherine Rawlinson, Kate Rylands, Mina Sheppard, Nathan Williams, Victoria Williams and Ella Wright. This work was funded by NERC standard grant NE/N010221/1. JH was supported by Czech Science Foundation grant no. 17-27184Y.

## References

Anderson, T.L. & Whiteman, H.H. (2015). Asymmetric effects of intra- and interspecific competition on a pond-breeding salamander. Ecology, 96, 1681 – 1690.

Arthur, A.L., Weeks, A.R. & Sgro, C.M. (2008). Investigating latitudinal clines for life history and stress resistance traits in Drosophila simulans from eastern Australia. Journal of Evolutionary Biology, 21, 1470 – 1479.

Barraclough, T.G. (2015). How do species interactions affect evolutionary dynamics across whole communities? Annual Review of Ecology, Evolution and Systematics 46, 25 – 48.

Bates, D., Machler, M., Bolker, B.M. & Walker, S.C. (2015). Fitting Linear Mixed-Effects Models Using lme4. Journal of Statistical Software, 67, 1 – 48.

Bischoff, A., Crémieux, L., Smilauerova, M., Lawson, C.S., Mortimer, S.R., Dolezal, J., Lanta, V., Edwards, A.R., Brook, A.J., Macel, M., Leps, J., Steinger, T. & Müller-Schärer, H. (2006). Detecting local adaptation in widespread grassland species -the importance of scale and local plant community. Journal of Ecology, 94, 1130 – 1142.

Bridle, J.R., Gavaz, S. & Kennington, W.J. (2009). Testing limits to adaptation along altitudinal gradients in rainforest Drosophila. Proceedings of the Royal Society B: Biological Sciences, 276, 1507 – 1515.

Bridle, J.R., Kawata, M. & Butlin, R.K. (2019). Local adaptation stops where ecological gradients steepen or are interrupted. Evolutionary Applications, 12, 1449 – 1462.

Coley, P.D. & Barone, J.A. (1996). Herbivory and plant defenses in tropical forests. Annual Review of Ecology and Systematics, 27, 305 – 335

Darwin, C.R. (1859). On the origins of species by means of natural selection, or the preservation of favoured races in the struggle for life. John Murray, London UK

Davis, A.J., Jenkinson, L.S., Lawton, J.H., Shorrocks, B. & Wood, S. (1998). Making mistakes when predicting shifts in species range in response to global warming. Nature, 391, 783 – 786.

Fiegna, F., Scheuerl, T., Moreno-Letelier, A., Bell, T. & Barraclough, T.G. (2015). Saturating effects of species diversity on life-history evolution in bacteria. Proceedings of the Royal Society of London B., 282, 20151794.

Godsoe, W., Murray, R. & Plank, M.J. (2015). The effect of competition on species’ distributions depends on coexistence, rather than scale alone. Ecography 38, 1071 – 1079.

Godsoe, W., Franklin, J. & Blanchet, F.G. (2017). Effects of biotic interactions on modeled species’ distribution can be masked by environmental gradients. Ecology and Evolution, 7, 654 – 664.

Grant, E.H.C., Brand, A.B., De Wekker, S.F.J., Lee, T.R. & Wofford, J.E.B. (2018). Evidence that climate sets the lower elevation range limit in a high-elevation endemic salamander. Ecology and Evolution, 8, 7553 – 7562.

Griffiths, J.A., Schiffer, M. & Hofmann, A.A. (2005). Clinal variation and laboratory adaptation in the rainforest species Drosophila birchii for stress resistance, wing size, wing shape and development time. Journal of Evolutionary Biology, 18, 213 – 222.

Hall, J.P.J., Harrison, E. & Brockhurst, M.A. (2018). Competitive species interactions constrain abiotic adaptation in a bacterial soil community. Evolution Letters, 2, 580 – 589.

Hargreaves, A.L., Germain, R.M., Bontrager, M., Persi, J. & Angert, A.L. (2020). Local adaptation to biotic interactions: a meta-analysis across latitudes. The American Naturalist 195, 395 – 411.

Inouye, B.D. (2001). Response surface experimental designs for investigating interspecific competition. Ecology, 82, 2696 – 2706.

Jeffs, C.T., Terry, J.C.D., Higgie, M., Jandovà, A., Konvickovà, H., Brown, J.J., Lue, C-H., Schiffer, M., O’Brien, E.K., Bridle, J.R., Hrček, J. & Lewis, O.T. (2020). Molecular analyses reveal consistent food web structure with elevation in rainforest *Drosophila*-parasitoid communities. BioRxiv DOI: https://doi.org/10.1101/2020.07.21.213678.

Jousset, A., Eisenhauer, N., Merker, M., Mouquet, N. & Scheu, S. (2016). High functional diversity stimulates diversification in experimental microbial communities. Science Advances, 2, e1600124.

Kawecki, T.J. & Ebert, D. (2004). Conceptual issues in local adaptation. Ecology Letters, 7, 1225 – 1241.

Körner, C. (2007). The use of ‘altitude’ in ecological research. Trends in Ecology & Evolution, 22, 569 – 574.

Lawrence, D., Fiegna, F., Behrends, V., Bundy, J.G., Phillimore, A.B., Bell, T. & Barraclough, T.G. (2012). Species interactions alter evolutionary responses to a novel environment. PLOS Biology, 10, e1001330.

Long, T.A.F. & Pischedda, A. (2005). Do female Drosophila melanogaster adaptively bias offspring sex ratios in relation to the age of their mate? Proceedings of the Royal Society B: Biological Sciences, 272, 1781 – 1787.

Lurgi, M., López, B.C. & Montoya, J.M. (2012). Novel communities from climate change. Philosophical Transactions of the Royal Society B, 367, 2913 – 2922.

Mange, A.P. (1970). Possible nonrandom utilization of X- and Y-bearing sperm in Drosophila melanogaster. Genetics, 65, 95 – 106.

McCabe, J. & Partridge, L. (1997). An interaction between environmental temperature and genetic variation for body size for the fitness of adult female Drosophila melanogaster. Evolution, 51, 1164 – 1174.

Moles, A.T. & Ollerton, J. (2016). Is the notion that species interactions are stronger and more specialized in the tropics a zombie idea? Biotropica, 48, 141 – 145.

Morris, R.J., Sinclair, F.H. & Burwell, C.J. (2015). Food web structure changes with elevation but not rainforest stratum. Ecography, 38, 792 – 802.

O’Brien, E.K., Higgie, M., Reynolds, A., Hoffmann, A.A. & Bridle, J.R. (2017). Testing for local adaptation and evolutionary potential along altitudinal gradients in rainforest Drosophila: beyond laboratory estimates. Global Change Biology, 23, 1847 – 1860.

Partridge, L. & Farquhar, M. (1983). Lifetime mating success of male fruitflies (Drosophila melanogaster) is related to their size. Animal Behaviour, 31, 871 – 877.

Pearson, R.G. & Dawson, T.P. (2003). Predicting the impacts of climate change on the distribution of species: are bioclimate envelope models useful? Global Ecology and Biogeography, 12, 361 – 371.

Poisot, T., Bever, J.D., Nemri, A., Thrall, P.H. & Hochberg, M.E. (2011). A conceptual framework for the evolution of ecological specialisation. Ecology Letters, 14, 841 – 851.

Rahbek, C. (1995). The elevational gradient of species richness - a uniform pattern. Ecography, 18, 200 – 205.

Rice, K.J. & Knapp, E.E. (2008). Effects of competition and life history stage on the expression of local adaptation in two native bunchgrasses. Restoration Ecology, 16, 12 – 23.

Santos, M., Ruiz, A., Quezada-Diaz, J.E., Barbadilla, A. & Fontdevila, A. (1992). The evolutionary history of Drosophila buzzatii. XX. Positive phenotypic covariance between field adult fitness components and body size. Journal of Evolutionary Biology, 5, 403 – 422.

Santos, M., Fowler, K. & Partridge, L. (1994). Gene-environment interaction for body size and larval density in Drosophila melanogaster: an investigation of effects on development time, thorax length and adult sex ratio. Heredity, 72, 515 – 521

Schemske, D.W., Mittelbach, G.G., Cornell, H.V., Sobel, J.M. & Roy, K. (2009). Is there a latitudinal gradient in the importance of biotic interactions? Annual Review of Ecology, Evolution, and Systematics, 40, 245 – 269.

Scheuerl, T., Hopkins, M., Nowell, R.W., Rivett, D.W., Barraclough, T.G. & Bell, T. (2020). Bacterial adaptation is constrained in complex communities. Nature Communications, 11, 1 – 8.

Schiffer, M. & McEvey, S.F. (2006). Drosophila bunnanda - a new species from northern Australia with notes on other Australian members of the montium subgroup (Diptera:Drosophilidae). Zootaxa, 1333, 1–23.

Stuart, Y.E., Campbell, T.S., Hohenlohe, P.A., Reynolds, R.G., Revell, L.J. & Losos, J.B. (2014). Rapid evolution of a native species following invasion by a congener. Science, 346, 463 – 466.

Siepielski, A.M., Nemirov, A., Cattivera, M. & Nickerson, A. (2016). Experimental evidence for an eco-evolutionary coupling between local adaptation and intraspecific competition. The American Naturalist 187, 447 – 456.

Trivers, R.L. & Willard, D.E. (1973). Natural selection of parental ability to vary sex ratio of offspring. Science, 179, 90 – 92.

van Heerwaarden, B., Kellermann, V., Schiffer, M., Blacket, M., Sgrò, C.M. & Hoffmann, A.A. (2009). Testing evolutionary hypotheses about species borders: patterns of genetic variation towards the southern borders of two rainforest Drosophila and a related habitat generalist. Proceedings of the Royal Society B., 276, 1517 – 1526.

Wisz, M.S., Pottier, J., Kissling, W.D., Pellissier, L., Lenoir, J., Damgaard, C.F. et al. (2013). The role of biotic interactions in shaping distributions and realised assemblages of species: implications for species distribution modelling. Biological Reviews, 88, 15–30.

