## Supplementary Information for "Fitness effects of competition within and between species change across species’ ranges, and reveal limited local adaptation in rainforest *Drosophila*"

**Supplementary Note 1: Establishment of isofemale lines and mass-bred populations of *D. birchii* and D. bunnanda used in the caged transplant experiment**

***Establishment of isofemale lines***

We placed field-mated females individually in 40 ml glass vials containing 10 ml of standard *Drosophila* medium (agar, raw sugar, inactive yeast, propionic acid and methyl-4-hydroxybenzoate), topped with a few grains of live yeast to stimulate egg-laying. We left them to oviposit for 3 – 4 days and then moved them to a fresh food vial. This process was repeated until they ceased laying eggs, and their offspring were left to emerge. We mixed all offspring of the same female together to found the next generation and establish the isofemale line. Lines were were maintained at 23 °C on a 12:12 hr light:dark cycle for 10 months (~20 generations) prior to the establishment of mass-bred populations.

***Establishment of mass-bred populations***

We established a mass-bred population for each of the sites where *D. birchii* were collected (eight populations in total: two high and two low elevation populations from each of Paluma and Mt Lewis) by mixing flies from the 10 isofemale lines from that site. We established a single mixed mass-bred population of *D. bunnanda* (a mixture of low elevation populations from Paluma and Mt Lewis) by mixing all 10 isofemale lines of this species.

To construct mass-bred populations, the 10 contributing lines were divided into two groups of 5 lines each (Group A and Group B for each population). Flies emerging from each line were sexed on the day of emergence and held separately by sex to ensure they were unmated. For each population, we initially set up 6 x 250 ml rearing bottles with 100 ml standard *Drosophila* media, with 100 flies per bottle. Three of these bottles contained 10 virgin males from Group A lines and 10 virgin females from Group B lines, while the remaining three bottles contained the reverse combination. Flies emerging across the six replicate bottles for a population were mixed each generation. We used this approach to try to preserve genetic variation within each population, as it ensured that all offspring emerging from the mass-bred population in each generation after establishment had alleles contributed by at least two isofemale lines. Mass-bred populations were maintained at 23 °C on a 12:12 hr light:dark cycle for two generations prior to being used in the field transplant experiment. The number of flies laying in each bottle was limited to 100 each generation to keep the density low and minimize variation in size or productivity due to larval competition. The number of replicate bottles per population was doubled to 12 in the generation preceding the experiment, to ensure sufficient flies were available in the experimental generation.

**Supplementary Note 2: Abiotic variation along gradient**

We placed a Carbon-51 USB data logger (Sensormetrix, UK) alongside experimental vials inside each of five randomly chosen holders at each transplant location (one per block), to measure temperature and humidity every 10 minutes during the transplant experiment. Elevation was a major source of variation in abiotic variables, explaining more than 60% of variation in temperature and around 30% of variation in humidity over the duration of the experiment (Figure S1). Mean daily ambient temperature was 21.4 °C at the low site and, respectively, 2.3 °C and 5.2 °C cooler at the mid and high sites (Figure S1). Humidity increased with elevation, from 84.8% at the low site to 95.2% and 96.9% at the mid and high sites (Figure S1). There was substantial variation among days in both measures (11.4% of variance in temperature, 39.4% of variance in humidity), but no significant variation among blocks within elevations (Figure S1).

**Supplementary Note 3: Pilot study to assess effect of intraspecific density on productivity in *Drosophila birchii***

We assessed the productivity of *D. birchii* in vials with varying numbers of adult flies, to determine the starting density at which intraspecific competition has a negative effect on fitness. We used the results of this experiment to select a range of total densities for our field transplant experiment that we were confident would be sufficiently high to induce competition within and between *D. birchii* and *D. bunnanda*.

We tested seven different starting densities of *D. birchii*: 4, 8, 16, 24, 32, 40, and 64 flies. In each case, we placed equal numbers of 7-day old unmated males and females in a 40ml vial with 5ml of standard *Drosophila* media (agar, sugar, inactive yeast, methyl hydroxybenzoate and propionic acid). We established 10 replicate vials of each starting density. We left flies to mate and lay for 7 days at 23 °C on a 12:12 hr light:dark cycle, then discarded them and added pupation card. We then left vials under these same conditions until all offspring had emerged. We counted the total number of offspring emerging from each vial and divided this by the number of laying females to give a measure of productivity.

There was no detectable decline in productivity with an increase in density from 4 flies to 8 flies, but above this density, productivity declined rapidly, such that the mean number of offspring per female at density = 24 was approximately one third of that when density = 8 (Figure S2). To capture this range, we chose to include total densities between 6 - 48 in our field transplant experiment (intraspecific densities of 6 – 24).

**Supplementary Note 4: Relevance of competition treatments used in the field transplant experiment to natural competition intensities**

Larval densities, and how they vary in natural populations are not known, making it challenging to relate densities used in this experiment to those experienced by free-living flies as larvae or as adults. However, larval density is a strong determinant of body size at a given developmental temperature in *Drosophila* (e.g. Miller & Thomas 1958; Santos et al. 1994). Comparison of the mean body size of field-caught flies at a site with that of the flies emerging from field vials at the same site was therefore used as an indicator of field larval density, and therefore of the ecological relevance of the density treatments used in the transplant cages. To make this comparison, we collected males and females of each species from each transplant elevation at Paluma (only low and middle elevations for *D. bunnanda*) over the same period as the caged transplant experiment was running. We measured wing size, as a proxy for body size, using the same method as for the experimental flies (Griffiths et al. 2005), for approximately 50 flies of each sex and species from each elevation (mean = 50.4, range = 8 – 135). The mean wing sizes of flies emerging from field cages were within one standard deviation of the wing size of field-caught flies for both sexes of both species at all competition densities at most transplant sites (Figure 3), providing good evidence that the competition treatments used in our transplant experiments are similar to the range of larval densities found in nature, across all of the elevations used.

**Supplementary Figures**

| Elevation | Mean daily min temperature (SD) |
| --- | --- |
| Low | 18.79 |
| Mid | 17.0188 |
| High | 14.373 |

| Elevation | Mean daily temperature (SD) |
| --- | --- |
| Low | 21.3599 |
| Mid | 19.0831 |
| High | 16.1461 |

| Elevation | Mean daily max temperature (SD) |
| --- | --- |
| Low | 26.8277 |
| Mid | 21.6788 |
| High | 18.22 |

(i) Temperature

| Elevation | Mean daily min % humidity (SD) |
| --- | --- |
| Low | 65.7938 |
| Mid | 84.3 |
| High | 90.003 |

| Elevation | Mean daily % humidity (SD) |
| --- | --- |
| Low | 84.85 |
| Mid | 95.1799 |
| High | 96.9188 |

| Elevation | Mean daily max % humidity (SD) |
| --- | --- |
| Low | 95.8077 |
| Mid | 99.4 |
| High | 99.617 |

(ii) Humidity
(A) Average daily minimum, mean and maximum (i) temperature and (ii) humidity at each transplant elevation. Error bars are standard deviations.

Low elevation

Temperature

| Day of experiment | Mean | Min | Max |
| --- | --- | --- | --- |
| 2 | 23.4656 | 19.8 | 36.8 |
| 3 | 22.8253 | 18.8 | 31.1 |
| 4 | 23.8781 | 20.6 | 29.8 |
| 5 | 20.8143 | 15.5 | 29.3 |
| 6 | 20.7869 | 16.3 | 28.3 |
| 7 | 21.2047 | 19.3 | 27.3 |
| 8 | 22.159 | 19.1 | 30 |
| 9 | 22.4086 | 19 | 30.5 |
| 10 | 22.4089 | 19 | 30.8 |
| 11 | 22.3285 | 19.6 | 27 |
| 12 | 20.4215 | 19.5 | 21.4 |
| 13 | 18.7706 | 17.7 | 19.8 |
| 14 | 18.312 | 17.7 | 22.4 |

Mid elevation

Temperature

| Day of experiment | Mean | Min | Max |
| --- | --- | --- | --- |
| 2 | 19.2675 | 16.3 | 22.5 |
| 3 | 19.5571 | 16.4 | 23.6 |
| 4 | 20.2901 | 17.7 | 22.9 |
| 5 | 17.629 | 14.9 | 21.6 |
| 6 | 18.1857 | 14.1 | 22.2 |
| 7 | 18.9429 | 17.8 | 20.7 |
| 8 | 19.2676 | 16.9 | 22.7 |
| 9 | 18.8546 | 16.8 | 23.2 |
| 10 | 19.1803 | 16.5 | 22.3 |
| 11 | 20.2503 | 18.7 | 23.6 |
| 12 | 19.0994 | 18.1 | 20 |
| 13 | 17.3678 | 16.4 | 18.4 |
| 14 | 18.371 | 16.2 | 21.6 |
| 15 | 19.6517 | 17.5 | 22.1 |
| 16 | 19.2689 | 16.6 | 23.3 |
| 17 | 20.1453 | 19.7 | 21.4 |

High elevation

Temperature

| Day of experiment | Mean | Min | Max |
| --- | --- | --- | --- |
| 2 | 16.151 | 13.4 | 20.2 |
| 3 | 16.7947 | 14 | 19.6 |
| 4 | 16.3922 | 13.9 | 19.4 |
| 5 | 14.1908 | 12 | 18.8 |
| 6 | 15.489 | 11.7 | 19.8 |
| 7 | 16.1389 | 15.1 | 17.6 |
| 8 | 16.6508 | 14.6 | 19.4 |
| 9 | 16.7617 | 14.5 | 21.1 |
| 10 | 16.9683 | 14.3 | 20.9 |
| 11 | 17.3074 | 16 | 18.7 |
| 12 | 16.5042 | 15.3 | 17.6 |
| 13 | 14.6824 | 13.5 | 15.5 |
| 14 | 15.9907 | 13.5 | 18.3 |
| 15 | 16.8006 | 15.3 | 18.6 |
| 16 | 16.8476 | 14 | 21 |
| 17 | 17.5319 | 16.8 | 18.8 |
| 18 | 16.5399 | 14.5 | 19.9 |
| 19 | 16.2017 | 14 | 20.6 |
| 20 | 15.5469 | 12.9 | 19.5 |
| 21 | 13.4307 | 12 | 15.9 |

(i) Temperature

Low elevation

% Humidity

| Day of experiment | Mean | Min | Max |
| --- | --- | --- | --- |
| 2 | 67.6763 | 33.3 | 89.7 |
| 3 | 77.7 | 52.3 | 91 |
| 4 | 70.6133 | 46.4 | 90.2 |
| 5 | 67.7662 | 35.6 | 85.9 |
| 6 | 82.536 | 53.5 | 100 |
| 7 | 93.7439 | 65.4 | 100 |
| 8 | 87.6669 | 57.7 | 100 |
| 9 | 82.7367 | 56.8 | 96.6 |
| 10 | 81.8938 | 48.5 | 97 |
| 11 | 90.8329 | 66.2 | 100 |
| 12 | 100 | 100 | 100 |
| 13 | 100 | 100 | 100 |
| 14 | 99.884 | 92 | 100 |

Mid elevation

% Humidity

| Day of experiment | Mean | Min | Max |
| --- | --- | --- | --- |
| 2 | 86.551 | 56.5 | 97.2 |
| 3 | 92.5356 | 77.9 | 99.3 |
| 4 | 86.5211 | 68.3 | 99.3 |
| 5 | 79.466 | 50.9 | 96.6 |
| 6 | 92.8897 | 76.5 | 100 |
| 7 | 99.9071 | 96.5 | 100 |
| 8 | 97.9082 | 81.9 | 100 |
| 9 | 97.0236 | 82.7 | 100 |
| 10 | 96.116 | 78.2 | 100 |
| 11 | 97.285 | 83.7 | 100 |
| 12 | 100 | 100 | 100 |
| 13 | 100 | 100 | 100 |
| 14 | 99.7537 | 92 | 100 |
| 15 | 98.9835 | 88.9 | 100 |
| 16 | 98.4574 | 82.6 | 100 |
| 17 | 99.4869 | 93.5 | 100 |

High elevation

% Humidity

| Day of experiment | Mean | Min | Max |
| --- | --- | --- | --- |
| 2 | 91.6521 | 59.7 | 100 |
| 3 | 95.8867 | 86.3 | 100 |
| 4 | 95.9961 | 76.1 | 100 |
| 5 | 90.2131 | 55.6 | 100 |
| 6 | 98.1021 | 86.9 | 100 |
| 7 | 100 | 100 | 100 |
| 8 | 99.9953 | 97.3 | 100 |
| 9 | 99.6465 | 89.8 | 100 |
| 10 | 99.3553 | 83 | 100 |
| 11 | 100 | 100 | 100 |
| 12 | 100 | 100 | 100 |
| 13 | 100 | 100 | 100 |
| 14 | 100 | 100 | 100 |
| 15 | 100 | 100 | 100 |
| 16 | 99.8718 | 92.4 | 100 |
| 17 | 100 | 100 | 100 |
| 18 | 99.2418 | 88 | 100 |
| 19 | 81.9004 | 63.8 | 98.7 |
| 20 | 91.1272 | 67.3 | 100 |
| 21 | 95.387 | 79.3 | 100 |

(ii) Humidity
(B) Daily mean, minimum and maximum (i) temperature and (ii) humidity at each transplant elevation. In each case, the daily mean is the solid line, the daily minimum is the lower dashed line, and the daily maximum is the upper dashed line.

| **Abiotic variable** | **Source of variation** | **Variance** | **% Variance** | **χ^2^ (1 d.f.)** | ***P*** |
| --- | --- | --- | --- | --- | --- |
| Temperature | Elevation | 7.234 | 61.58 | 52.99 | 3.36 x 10^-13^ |
|  | Block (Elevation) | 0.000 | 0.00 | 0.00 | 1 |
|  | Day(Block(Elevation)) | 1.336 | 11.37 | 9255.5 | <2.2 x 10^-16^ |
|  | Residual | 3.178 | 27.05 |  |  |
| Humidity | Elevation | 41.83 | 29.36 | 29.89 | 4.57 x 10^-8^ |
|  | Block (Elevation) | 0.000 | 0.00 | 0.00 | 1 |
|  | Day(Block(Elevation)) | 56.08 | 39.36 | 25614 | <2.2 x 10^-16^ |
|  | Residual | 44.57 | 31.28 |  |  |

(C) Spatial and temporal variance in abiotic variables (temperature and humidity) across the Paluma elevation gradient used for the caged transplant experiment. Shown are variances and % variances among transplant elevations, blocks within transplant elevations, and days, estimated from linear mixed models with each of these variables included as a random effect. The significance of each source of variation was estimated by comparing the log-likelihood of the full model with a model where that factor was excluded, using a χ^2^ test with one degree of freedom. Residuals indicate daily variation within blocks at each elevation.

**Figure S1.**

No. offspring per D. birchii female (±SE)

| Number of D. birchii | Offspring per female |
| --- | --- |
| 4 | 23.05 |
| 8 | 23.625 |
| 16 | 14.85 |
| 24 | 8.325 |
| 32 | 8.53125 |
| 40 | 7.115 |
| 64 | 4 |

**Figure S2.** Mean productivity (number of offspring per female) of *Drosophila birchii* reared in the laboratory at seven different densities between 4 and 64 flies (equal numbers of males and females in each). There were 10 replicate vials per density treatment, and the error bars indicate standard errors among replicates.

(B) Predicted relative abundance assuming no interspecific competition and constant intraspecific density (density = 12)
(A) Observed relative abundance at field sites

Observed relative abundance

| Elevation | D. birchii | D. bunnanda |
| --- | --- | --- |
| Low | 1 | 2.51812 |
| Mid | 4.61775 | 4.53623 |
| High | 17.663 | 0 |

(C) Predicted relative abundance assuming no interspecific competition and field-observed intraspecific density

Predicted relative abundance

| Elevation | D. birchii | D. bunnanda |
| --- | --- | --- |
| Low | 1 | 0.921659 |
| Mid | 0.528418 | 0.427035 |
| High | 0.190476 | 0.394777 |

Predicted relative abundance

| Elevation | D. birchii | D. bunnanda |
| --- | --- | --- |
| Low | 1 | 1.59612 |
| Mid | 1.89319 | 1.22922 |
| High | 0.920289 | 0 |

(D) Predicted relative abundance assuming field-observed intraspecific and interspecific density

Predicted relative abundance

| Elevation | D. birchii | D. bunnanda |
| --- | --- | --- |
| Low | 1 | 1.79568 |
| Mid | 1.53056 | 1.11169 |
| High | 1.08145 | 0 |

**Figure S4:** Figures comparing predicted relative abundances of *D. birchii* and *D. bunnanda* based on competition effects on productivity in the field transplant experiment with observed relative abundances in field populations. (A) Observed relative abundance of *Drosophila birchii* and *D. bunnanda* based on population sampling at the low, mid and high elevation field sites where the caged transplant experiment was conducted. The remaining panels show predicted relative abundances of each species at each elevation based on their productivity in field cages and effects of intra and interspecific competition estimated in elevation-specific models from our experiment. Shown are three different sets of predictions based on different assumptions about the type and frequency of competitive interactions: (B) Predicted relative abundance assuming no interspecific competition and the same intermediate level of intraspecific competition in both species at all sites (using productivity with intraspecific density = 12, interspecific density = 0 in cages), (C) Predicted relative abundance assuming no interspecific competition and intraspecific interactions at a frequency proportional to field observations of the abundance of each species, and (D) Predicted relative abundance assuming intraspecific and interspecific interactions at a frequency proportional to the field abundance of each species. In all cases, observed/predicted relative abundance was calculated relative to the abundance of *D. birchii* at the low elevation site. Note that the predicted abundance of *D. bunnanda* at the high elevation site is 0 in (C) and (D) because per female productivity was multiplied by the observed field abundance to calculate predicted relative abundance, and *D. bunnanda* was not found at this site.

**Supplementary tables**

**Table S1.** Results of tests of effects of inter and intraspecific density and source elevation (*D. birchii* only), and all two-way interactions on (A) productivity, (B) offspring sex ratio, (C) emergence time, and (D) wing size of *Drosophila birchii* and *D. bunnanda* at low, mid and high elevation transplant sites. These were obtained from (generalised) linear mixed models fitted separately for each species, trait and elevation, and separately for males and females for emergence time and wing size. The significance of each factor was evaluated using a χ2 to compare the log-likelihood of models with and without the term included. *P*-values for fixed effects were obtained using the same method of log-likelihood comparison as described in Table 1. Within each species and site, we used a Bonferroni-corrected significance threshold of *P* = 0.013 to test each fixed effect, to account for testing of multiple traits. Tests where the *P*-value for a trait was below this threshold are highlighted and in bold italics. Tests where *P* < 0.05 but not below the corrected threshold are in italics.

| (A) **Productivity** | | | | | |
| --- | --- | --- | --- | --- | --- |
| **Species** | **Elevation** | **Fixed effect** | **χ^2^_(1 df)_** | | ***P*** |
| ***Drosophila birchii*** | **Low** | ***Intraspecific density*** | ***124.4*** | | ***<2.2 x 10^-16^*** |
|  |  | ***Interspecific density*** | ***128.7*** | | ***<2.2 x 10^-16^*** |
|  |  | Source elevation | 1.778 | | 0.182 |
|  |  | ***Intraspecific x interspecific density*** | ***41.15*** | | ***1.41 x 10^-10^*** |
|  |  | Intraspecific density x Source elevation | 1.751 | | 0.186 |
|  |  | Interspecific density x Source elevation | 0.165 | | 0.685 |
|  | **Mid** | ***Intraspecific density*** | ***98.62*** | | ***<2.2 x 10^-16^*** |
|  |  | ***Interspecific density*** | ***123.4*** | | ***<2.2 x 10^-16^*** |
|  |  | Source elevation | 0.185 | | 0.668 |
|  |  | ***Intraspecific x interspecific density*** | ***63.29*** | | ***1.73 x 10^-15^*** |
|  |  | Intraspecific density x Source elevation | 1.153 | | 0.283 |
|  |  | Interspecific density x Source elevation | 0.786 | | 0.375 |
|  | **High** | ***Intraspecific density*** | ***83.67*** | | ***<2.2 x 10^-16^*** |
|  |  | ***Interspecific density*** | ***58.71*** | | ***1.83 x 10^-14^*** |
|  |  | Source elevation | 0.607 | | 0.436 |
|  |  | ***Intraspecific x interspecific density*** | ***36.51*** | | ***1.52 x 10^-9^*** |
|  |  | Intraspecific density x Source elevation | 0.014 | | 0.905 |
|  |  | Interspecific density x Source elevation | 2.381 | | 0.123 |
| ***Drosophila bunnanda*** | **Low** | ***Intraspecific density*** | ***88.37*** | | ***<2.2 x 10^-16^*** |
|  |  | ***Interspecific density*** | ***35.45*** | | ***2.62 x 10^-9^*** |
|  |  | ***Intraspecific x interspecific density*** | ***10.92*** | | ***9.53 x 10^-4^*** |
|  | **Mid** | ***Intraspecific density*** | ***64.09*** | | ***1.19 x 10^-15^*** |
|  |  | ***Interspecific density*** | ***41.02*** | | ***1.51 x 10^-10^*** |
|  |  | ***Intraspecific x interspecific density*** | ***13.08*** | | ***2.99 x 10^-4^*** |
|  | **High** | ***Intraspecific density*** | ***24.20*** | | ***8.71 x 10^-7^*** |
|  |  | ***Interspecific density*** | ***34.15*** | | ***5.12 x 10^-9^*** |
|  |  | ***Intraspecific x interspecific density*** | ***17.49*** | | ***2.90 x 10^-5^*** |
| (B) **Offspring sex ratio** | | | | | |
| **Species** | **Elevation** | **Fixed effect** | **χ^2^_(1 df)_** | | ***P*** |
| ***Drosophila birchii*** | **Low** | Intraspecific density | 0.048 | | 0.827 |
|  |  | ***Interspecific density*** | ***26.32*** | | ***2.89 x 10^-7^*** |
|  |  | Source elevation | 0.071 | | 0.791 |
|  |  | Intraspecific x interspecific density | 0.638 | | 0.424 |
|  |  | Intraspecific density x Source elevation | 0.000 | | 0.983 |
|  |  | Interspecific density x Source elevation | 0.008 | | 0.929 |
|  | **Mid** | Intraspecific density | 0.043 | | 0.837 |
|  |  | Interspecific density | 1.782 | | 0.182 |
|  |  | Source elevation | 1.561 | | 0.212 |
|  |  | Intraspecific x interspecific density | 0.004 | | 0.952 |
|  |  | Intraspecific density x Source elevation | 0.102 | | 0.750 |
|  |  | Interspecific density x Source elevation | 0.066 | | 0.798 |
|  | **High** | Intraspecific density | 0.434 | | 0.510 |
|  |  | Interspecific density | 0.116 | | 0.734 |
|  |  | Source elevation | 0.003 | | 0.955 |
|  |  | Intraspecific x interspecific density | 0.241 | | 0.624 |
|  |  | Intraspecific density x Source elevation | 1.179 | | 0.278 |
|  |  | Interspecific density x Source elevation | 0.016 | | 0.899 |
| ***Drosophila bunnanda*** | **Low** | ***Intraspecific density*** | ***10.07*** | | ***0.002*** |
|  |  | *Interspecific density* | *5.034* | | *0.025* |
|  |  | Intraspecific x interspecific density | 0.000 | | 0.985 |
|  | **Mid** | Intraspecific density | 1.580 | | 0.209 |
|  |  | Interspecific density | 0.622 | | 0.430 |
|  |  | Intraspecific x interspecific density | 1.404 | | 0.236 |
|  | **High** | Intraspecific density | 0.065 | | 0.800 |
|  |  | Interspecific density | 0.302 | | 0.583 |
|  |  | Intraspecific x interspecific density | 1.868 | | 0.172 |

| (C) **Emergence time** | | | | | |
| --- | --- | --- | --- | --- | --- |
| **Species** | **Elevation** | **Sex** | **Fixed effect** | **χ^2^_(1 df)_** | ***P*** |
| ***Drosophila birchii*** | **Low** | **Male** | ***Intraspecific density*** | ***17.32*** | ***3.16 x 10^-5^*** |
|  |  |  | ***Interspecific density*** | ***8.593*** | ***0.003*** |
|  |  |  | Source elevation | 2.226 | 0.136 |
|  |  |  | Intra x Inter density | 0.078 | 0.780 |
|  |  |  | Intra density x Source elevation | 0.385 | 0.535 |
|  |  |  | Inter density x Source elevation | 0.000 | 0.987 |
|  |  | **Female** | ***Intraspecific density*** | ***6.410*** | ***0.011*** |
|  |  |  | ***Interspecific density*** | ***13.19*** | ***2.81 x 10^-4^*** |
|  |  |  | Source elevation | 3.370 | 0.066 |
|  |  |  | *Intra x Inter density* | *5.979* | *0.014* |
|  |  |  | *Intra density x Source elevation* | *5.641* | *0.018* |
|  |  |  | Inter density x Source elevation | 0.032 | 0.857 |
|  | **Mid** | **Male** | Intraspecific density | 3.590 | 0.058 |
|  |  |  | *Interspecific density* | *4.173* | *0.041* |
|  |  |  | Source elevation | 0.065 | 0.799 |
|  |  |  | Intra x Inter density | 0.000 | 0.977 |
|  |  |  | Intra density x Source elevation | 1.106 | 0.293 |
|  |  |  | Inter density x Source elevation | 0.022 | 0.881 |
|  |  | **Female** | Intraspecific density | 2.749 | 0.097 |
|  |  |  | ***Interspecific density*** | ***9.239*** | ***0.002*** |
|  |  |  | Source elevation | 0.720 | 0.396 |
|  |  |  | ***Intra x Inter density*** | ***6.545*** | ***0.011*** |
|  |  |  | ***Intra density x Source elevation*** | ***7.482*** | ***0.006*** |
|  |  |  | Inter density x Source elevation | 0.043 | 0.836 |
|  | **High** | **Male** | Intraspecific density | 0.629 | 0.428 |
|  |  |  | ***Interspecific density*** | ***8.858*** | ***0.003*** |
|  |  |  | Source elevation | 0.344 | 0.557 |
|  |  |  | Intra x Inter density | 0.155 | 0.694 |
|  |  |  | Intra density x Source elevation | 0.643 | 0.423 |
|  |  |  | Inter density x Source elevation | 0.552 | 0.457 |
|  |  | **Female** | Intraspecific density | 0.333 | 0.564 |
|  |  |  | ***Interspecific density*** | ***10.17*** | ***0.001*** |
|  |  |  | Source elevation | 0.325 | 0.568 |
|  |  |  | Intra x Inter density | 0.059 | 0.808 |
|  |  |  | Intra density x Source elevation | 0.039 | 0.843 |
|  |  |  | Inter density x Source elevation | 0.178 | 0.673 |
| ***Drosophila bunnanda*** | **Low** | **Male** | ***Intraspecific density*** | ***9.405*** | ***0.002*** |
|  |  |  | Interspecific density | 0.116 | 0.734 |
|  |  |  | Intra x Inter density | 0.786 | 0.375 |
|  |  | **Female** | ***Intraspecific density*** | ***6.355*** | ***0.012*** |
|  |  |  | *Interspecific density* | *5.317* | *0.021* |
|  |  |  | Intra x Inter density | 1.226 | 0.268 |
|  | **Mid** | **Male** | Intraspecific density | 0.082 | 0.775 |
|  |  |  | Interspecific density | 0.113 | 0.737 |
|  |  |  | Intra x Inter density | 0.879 | 0.348 |
|  |  | **Female** | Intraspecific density | 0.086 | 0.769 |
|  |  |  | Interspecific density | 0.042 | 0.839 |
|  |  |  | Intra x Inter density | 0.446 | 0.504 |
|  | **High** | **Male** | Intraspecific density | 0.000 | 0.989 |
|  |  |  | Interspecific density | 0.175 | 0.676 |
|  |  |  | *Intra x Inter density* | *4.067* | *0.044* |
|  |  | **Female** | Intraspecific density | 0.040 | 0.842 |
|  |  |  | Interspecific density | 0.369 | 0.544 |
|  |  |  | Intra x Inter density | 0.002 | 0.965 |
| (D) **Wing size** | | | | | |
| **Species** | **Elevation** | **Sex** | **Fixed effect** | **χ^2^_(1 df)_** | ***P*** |
| ***Drosophila birchii*** | **Low** | **Male** | ***Intraspecific density*** | ***78.04*** | ***<2.2 x 10^-16^*** |
|  |  |  | ***Interspecific density*** | ***20.26*** | ***6.76 x 10^-6^*** |
|  |  |  | Source elevation | 0.757 | 0.384 |
|  |  |  | ***Intra x Inter density*** | ***8.459*** | ***0.004*** |
|  |  |  | Intra density x Source elevation | 0.154 | 0.695 |
|  |  |  | Inter density x Source elevation | 2.571 | 0.109 |
|  |  | **Female** | ***Intraspecific density*** | ***37.05*** | ***1.15 x 10^-9^*** |
|  |  |  | Interspecific density | 0.002 | 0.968 |
|  |  |  | Source elevation | 0.690 | 0.406 |
|  |  |  | *Intra x Inter density* | *5.431* | *0.020* |
|  |  |  | Intra density x Source elevation | 0.174 | 0.677 |
|  |  |  | Inter density x Source elevation | 0.424 | 0.515 |
|  | **Mid** | **Male** | ***Intraspecific density*** | ***28.67*** | ***8.58 x 10^-8^*** |
|  |  |  | ***Interspecific density*** | ***11.21*** | ***8.15 x 10^-4^*** |
|  |  |  | Source elevation | 0.562 | 0.453 |
|  |  |  | Intra x Inter density | 1.597 | 0.206 |
|  |  |  | Intra density x Source elevation | 0.075 | 0.784 |
|  |  |  | Inter density x Source elevation | 0.237 | 0.627 |
|  |  | **Female** | ***Intraspecific density*** | ***20.72*** | ***5.33 x 10^-6^*** |
|  |  |  | ***Interspecific density*** | ***12.01*** | ***5.30 x 10^-4^*** |
|  |  |  | *Source elevation* | *4.753* | *0.029* |
|  |  |  | ***Intra x Inter density*** | ***7.826*** | ***0.005*** |
|  |  |  | Intra density x Source elevation | 0.337 | 0.562 |
|  |  |  | Inter density x Source elevation | 0.358 | 0.550 |
|  | **High** | **Male** | ***Intraspecific density*** | ***20.39*** | ***6.33 x 10^-6^*** |
|  |  |  | *Interspecific density* | *6.215* | *0.013* |
|  |  |  | Source elevation | 1.403 | 0.236 |
|  |  |  | Intra x Inter density | 0.015 | 0.904 |
|  |  |  | *Intra density x Source elevation* | *4.402* | *0.036* |
|  |  |  | Inter density x Source elevation | 0.016 | 0.899 |
|  |  | **Female** | ***Intraspecific density*** | ***14.75*** | ***1.23 x 10^-4^*** |
|  |  |  | ***Interspecific density*** | ***13.71*** | ***2.14 x 10^-4^*** |
|  |  |  | Source elevation | 0.396 | 0.529 |
|  |  |  | Intra x Inter density | 2.046 | 0.153 |
|  |  |  | ***Intra density x Source elevation*** | ***8.421*** | ***0.004*** |
|  |  |  | Inter density x Source elevation | 2.159 | 0.142 |
| ***Drosophila bunnanda*** | **Low** | **Male** | ***Intraspecific density*** | ***14.99*** | ***1.08 x 10^-4^*** |
|  |  |  | ***Interspecific density*** | ***11.87*** | ***5.70 x 10^-4^*** |
|  |  |  | Intra x Inter density | 0.008 | 0.931 |
|  |  | **Female** | ***Intraspecific density*** | ***34.73*** | ***3.80 x 10^-9^*** |
|  |  |  | ***Interspecific density*** | ***12.14*** | ***4.94 x 10^-4^*** |
|  |  |  | Intra x Inter density | 1.114 | 0.291 |
|  | **Mid** | **Male** | ***Intraspecific density*** | ***8.462*** | ***3.63 x 10^-3^*** |
|  |  |  | Interspecific density | 2.993 | 0.084 |
|  |  |  | Intra x Inter density | 1.549 | 0.213 |
|  |  | **Female** | ***Intraspecific density*** | ***20.47*** | ***6.06 x 10^-6^*** |
|  |  |  | Interspecific density | 0.328 | 0.567 |
|  |  |  | ***Intra x Inter density*** | ***6.692*** | ***9.69 x 10^-3^*** |
|  | **High** | **Male** | Intraspecific density | 2.237 | 0.135 |
|  |  |  | ***Interspecific density*** | ***26.46*** | ***2.70 x 10^-7^*** |
|  |  |  | *Intra x Inter density* | *4.514* | *0.034* |
|  |  | **Female** | Intraspecific density | 0.222 | 0.638 |
|  |  |  | *Interspecific density* | *5.970* | *0.015* |
|  |  |  | *Intra x Inter density* | *5.676* | *0.017* |

**Table S2:** Heat map tables showing means (with standard deviations in brackets) of (A) productivity, (B) sex ratio, (C) emergence time, and (D) wing size at each density treatment at each transplant elevation for flies emerging from field vials for *Drosophila birchii* (left) and *D. bunnanda* (right). For each trait, cells are coloured from white – red, with darker shading corresponding to increasing mean values of the trait, to allow visual comparison among densities, elevations, species, and (for emergence time and wing size) between the sexes. Mean (SD) of each trait for each elevation, species and sex (where relevant) are shown in the top left corner of each table.

***Drosophila birchii***

***Drosophila bunnanda***

| **LOW**  5.52(5.25) | | **Density *D. bunnanda*** | | | | |
| --- | --- | --- | --- | --- | --- | --- |
|  |  | **0** | **6** | **12** | **18** | **24** |
| **Density *D. birchii*** | **0** |  |  |  |  |  |
|  | **6** | 15.87 (5.57) | 6.15 (2.85) |  | 3.91 (3.84) |  |
|  | **12** | 6.51 (2.16) |  | 2.55 (1.38) |  |  |
|  | **18** |  | 2.74 (1.07) |  |  |  |
|  | **24** | 3.39 (1.50) |  |  |  | 0.92 (0.82) |

| **MID**  2.44(2.82) | | **Density *D. bunnanda*** | | | | |
| --- | --- | --- | --- | --- | --- | --- |
|  |  | **0** | **6** | **12** | **18** | **24** |
| **Density *D. birchii*** | **0** |  |  |  |  |  |
|  | **6** | 7.47 (3.80) | 3.05 (2.07) |  | 1.17 (1.02) |  |
|  | **12** | 3.44 (1.56) |  | 1.00 (0.73) |  |  |
|  | **18** |  | 0.91 (0.54) |  |  |  |
|  | **24** | 1.14 (0.65) |  |  |  | 0.30 (0.26) |

| **MID**  1.49(1.34) | | **Density *D. bunnanda*** | | | | |
| --- | --- | --- | --- | --- | --- | --- |
|  |  | **0** | **6** | **12** | **18** | **24** |
| **Density *D. birchii*** | **0** |  |  | 2.78 (1.00) |  | 0.91 (0.47) |
|  | **6** |  | 2.97 (1.60) |  | 1.07 (0.54) |  |
|  | **12** |  |  | 0.98 (0.55) |  |  |
|  | **18** |  | 1.51 (1.30) |  |  |  |
|  | **24** |  |  |  |  | 0.24 (0.18) |

| **HIGH**  1.35(2.02) | | **Density *D. bunnanda*** | | | | |
| --- | --- | --- | --- | --- | --- | --- |
|  |  | **0** | **6** | **12** | **18** | **24** |
| **Density *D. birchii*** | **0** |  |  |  |  |  |
|  | **6** | 4.90 (3.22) | 1.66 (1.24) |  | 0.83 (0.89) |  |
|  | **12** | 1.24 (1.17) |  | 0.60 (0.61) |  |  |
|  | **18** |  | 0.38 (0.39) |  |  |  |
|  | **24** | 0.46 (0.30) |  |  |  | 0.16 (0.13) |

| **HIGH**  0.93(1.05) | | **Density *D. bunnanda*** | | | | |
| --- | --- | --- | --- | --- | --- | --- |
|  |  | **0** | **6** | **12** | **18** | **24** |
| **Density *D. birchii*** | **0** |  |  | 2.57 (1.28) |  | 0.88 (0.57) |
|  | **6** |  | 1.82 (1.65) |  | 0.64 (0.41) |  |
|  | **12** |  |  | 0.65 (0.44) |  |  |
|  | **18** |  | 0.67 (0.56) |  |  |  |
|  | **24** |  |  |  |  | 0.22 (0.20) |

(A) **Productivity (number of offspring per female)**

***Drosophila birchii***

***Drosophila bunnanda***

| **LOW**  0.51(0.15) | | **Density *D. bunnanda*** | | | | |
| --- | --- | --- | --- | --- | --- | --- |
|  |  | **0** | **6** | **12** | **18** | **24** |
| **Density *D. birchii*** | **0** |  |  |  |  |  |
|  | **6** | 0.45 (0.08) | 0.51 (0.13) |  | 0.55 (0.19) |  |
|  | **12** | 0.48 (0.09) |  | 0.55 (0.17) |  |  |
|  | **18** |  | 0.56 (0.14) |  |  |  |
|  | **24** | 0.45 (0.09) |  |  |  | 0.53 (0.24) |

| **LOW**  0.40(0.18) | | **Density *D. bunnanda*** | | | | |
| --- | --- | --- | --- | --- | --- | --- |
|  |  | **0** | **6** | **12** | **18** | **24** |
| **Density *D. birchii*** | **0** |  |  | 0.41 (0.08) |  | 0.47 (0.10) |
|  | **6** |  | 0.40 (0.13) |  | 0.50 (0.12) |  |
|  | **12** |  |  | 0.39 (0.19) |  |  |
|  | **18** |  | 0.30 (0.18) |  |  |  |
|  | **24** |  |  |  |  | 0.44 (0.27) |

| **MID**  0.49(0.25) | | **Density *D. bunnanda*** | | | | |
| --- | --- | --- | --- | --- | --- | --- |
|  |  | **0** | **6** | **12** | **18** | **24** |
| **Density *D. birchii*** | **0** |  |  |  |  |  |
|  | **6** | 0.48 (0.13) | 0.51 (0.27) |  | 0.43 (0.36) |  |
|  | **12** | 0.49 (0.13) |  | 0.55 (0.29) |  |  |
|  | **18** |  | 0.51 (0.26) |  |  |  |
|  | **24** | 0.45 (0.13) |  |  |  | 0.47 (0.33) |

| **MID**  0.45(0.27) | | **Density *D. bunnanda*** | | | | |
| --- | --- | --- | --- | --- | --- | --- |
|  |  | **0** | **6** | **12** | **18** | **24** |
| **Density *D. birchii*** | **0** |  |  | 0.54 (0.12) |  | 0.43 (0.17) |
|  | **6** |  | 0.41 (0.21) |  | 0.52 (0.20) |  |
|  | **12** |  |  | 0.48 (0.28) |  |  |
|  | **18** |  | 0.39 (0.31) |  |  |  |
|  | **24** |  |  |  |  | 0.46 (0.41) |

| **HIGH**  0.40(0.32) | | **Density *D. bunnanda*** | | | | |
| --- | --- | --- | --- | --- | --- | --- |
|  |  | **0** | **6** | **12** | **18** | **24** |
| **Density *D. birchii*** | **0** |  |  |  |  |  |
|  | **6** | 0.42 (0.20) | 0.38 (0.27) |  | 0.29 (0.37) |  |
|  | **12** | 0.40 (0.25) |  | 0.41 (0.34) |  |  |
|  | **18** |  | 0.48 (0.39) |  |  |  |
|  | **24** | 0.45 (0.30) |  |  |  | 0.32 (0.38) |

| **HIGH**  0.43(0.32) | | **Density *D. bunnanda*** | | | | |
| --- | --- | --- | --- | --- | --- | --- |
|  |  | **0** | **6** | **12** | **18** | **24** |
| **Density *D. birchii*** | **0** |  |  | 0.38 (0.14) |  | 0.35 (0.18) |
|  | **6** |  | 0.48 (0.25) |  | 0.47 (0.29) |  |
|  | **12** |  |  | 0.48 (0.31) |  |  |
|  | **18** |  | 0.32 (0.40) |  |  |  |
|  | **24** |  |  |  |  | 0.43 (0.38) |

(B) **Sex ratio (proportion of offspring male)**

***Drosophila birchii***

***Drosophila bunnanda***

| **LOW**  25.69  (6.36) | | **Density *D. bunnanda*** | | | | |
| --- | --- | --- | --- | --- | --- | --- |
|  |  | **0** | **6** | **12** | **18** | **24** |
| **Density *D. birchii*** | **0** |  |  |  |  |  |
|  | **6** | 26.59 (6.03) | 26.64 (6.30) |  | 25.25 (6.55) |  |
|  | **12** | 26.35 (6.47) |  | 25.76 (6.51) |  |  |
|  | **18** |  | 25.35 (6.39) |  |  |  |
|  | **24** | 24.33 (6.23) |  |  |  | 22.87 (5.65) |

| **LOW**  25.19 (6.13) | | **Density *D. bunnanda*** | | | | |
| --- | --- | --- | --- | --- | --- | --- |
|  |  | **0** | **6** | **12** | **18** | **24** |
| **Density *D. birchii*** | **0** |  |  | 25.60 (6.30) |  | 23.81 (6.39) |
|  | **6** |  | 26.33 (5.89) |  | 24.03 (6.18) |  |
|  | **12** |  |  | 25.46 (5.84) |  |  |
|  | **18** |  | 25.94 (6.43) |  |  |  |
|  | **24** |  |  |  |  | 24.83 (5.32) |

| **MID**  29.37  (6.34) | | **Density *D. bunnanda*** | | | | |
| --- | --- | --- | --- | --- | --- | --- |
|  |  | **0** | **6** | **12** | **18** | **24** |
| **Density *D. birchii*** | **0** |  |  |  |  |  |
|  | **6** | 29.81 (6.43) | 29.13 (6.41) |  | 30.45 (5.97) |  |
|  | **12** | 30.15 (6.07) |  | 27.33 (6.27) |  |  |
|  | **18** |  | 29.09 (6.28) |  |  |  |
|  | **24** | 28.59 (6.59) |  |  |  | 27.83 (5.75) |

| **MID**  29.54  (4.82) | | **Density *D. bunnanda*** | | | | |
| --- | --- | --- | --- | --- | --- | --- |
|  |  | **0** | **6** | **12** | **18** | **24** |
| **Density *D. birchii*** | **0** |  |  | 30.52 (4.20) |  | 29.27 (4.49) |
|  | **6** |  | 29.25 (5.13) |  | 29.05 (4.99) |  |
|  | **12** |  |  | 30.03 (4.49) |  |  |
|  | **18** |  | 29.17 (5.01) |  |  |  |
|  | **24** |  |  |  |  | 30.24 (4.87) |

| **HIGH**  35.25  (4.75) | | **Density *D. bunnanda*** | | | | |
| --- | --- | --- | --- | --- | --- | --- |
|  |  | **0** | **6** | **12** | **18** | **24** |
| **Density *D. birchii*** | **0** |  |  |  |  |  |
|  | **6** | 35.42 (4.53) | 34.04 (5.08) |  | 34.59 (5.83) |  |
|  | **12** | 36.50 (4.18) |  | 33.58 (5.18) |  |  |
|  | **18** |  | 34.60 (4.42) |  |  |  |
|  | **24** | 36.47 (4.42) |  |  |  | 34.22 (5.11) |

| **HIGH**  36.85  (3.76) | | **Density *D. bunnanda*** | | | | |
| --- | --- | --- | --- | --- | --- | --- |
|  |  | **0** | **6** | **12** | **18** | **24** |
| **Density *D. birchii*** | **0** |  |  | 38.09 (2.49) |  | 35.82 (4.69) |
|  | **6** |  | 36.69 (3.82) |  | 37.00 (3.32) |  |
|  | **12** |  |  | 36.23 (4.11) |  |  |
|  | **18** |  | 36.36 (4.23) |  |  |  |
|  | **24** |  |  |  |  | 37.57 (4.11) |

(C) **(i) Emergence time (days) males**

***Drosophila birchii***

***Drosophila bunnanda***

| **LOW**  24.88  (6.38) | | **Density *D. bunnanda*** | | | | |
| --- | --- | --- | --- | --- | --- | --- |
|  |  | **0** | **6** | **12** | **18** | **24** |
| **Density *D. birchii*** | **0** |  |  |  |  |  |
|  | **6** | 26.13 (5.99) | 25.30 (6.38) |  | 22.84 (6.10) |  |
|  | **12** | 25.52 (6.52) |  | 24.82 (6.91) |  |  |
|  | **18** |  | 23.67 (6.39) |  |  |  |
|  | **24** | 23.90 (6.25) |  |  |  | 23.55 (6.12) |

| **LOW**  25.42  (6.16) | | **Density *D. bunnanda*** | | | | |
| --- | --- | --- | --- | --- | --- | --- |
|  |  | **0** | **6** | **12** | **18** | **24** |
| **Density *D. birchii*** | **0** |  |  | 25.71 (6.07) |  | 24.04 (6.46) |
|  | **6** |  | 25.95 (6.01) |  | 24.14 (6.42) |  |
|  | **12** |  |  | 25.11 (6.12) |  |  |
|  | **18** |  | 26.69 (5.85) |  |  |  |
|  | **24** |  |  |  |  | 26.12 (5.58) |

| **MID**  29.10  (6.18) | | **Density *D. bunnanda*** | | | | |
| --- | --- | --- | --- | --- | --- | --- |
|  |  | **0** | **6** | **12** | **18** | **24** |
| **Density *D. birchii*** | **0** |  |  |  |  |  |
|  | **6** | 29.70 (6.26) | 29.31 (6.35) |  | 26.93 (6.53) |  |
|  | **12** | 29.84 (5.84) |  | 27.00 (6.39) |  |  |
|  | **18** |  | 29.57 (5.65) |  |  |  |
|  | **24** | 28.13 (6.15) |  |  |  | 28.27 (5.83) |

| **MID**  29.16  (5.12) | | **Density *D. bunnanda*** | | | | |
| --- | --- | --- | --- | --- | --- | --- |
|  |  | **0** | **6** | **12** | **18** | **24** |
| **Density *D. birchii*** | **0** |  |  | 30.08 (4.85) |  | 30.35 (4.20) |
|  | **6** |  | 28.80 (5.35) |  | 28.76 (5.14) |  |
|  | **12** |  |  | 29.49 (5.44) |  |  |
|  | **18** |  | 28.70 (4.99) |  |  |  |
|  | **24** |  |  |  |  | 29.06 (4.54) |

| **HIGH**  35.24 (4.94) | | **Density *D. bunnanda*** | | | | |
| --- | --- | --- | --- | --- | --- | --- |
|  |  | **0** | **6** | **12** | **18** | **24** |
| **Density *D. birchii*** | **0** |  |  |  |  |  |
|  | **6** | 35.55 (4.57) | 34.65 (4.97) |  | 34.53 (5.87) |  |
|  | **12** | 36.06 (4.92) |  | 34.14 (5.23) |  |  |
|  | **18** |  | 34.88 (5.12) |  |  |  |
|  | **24** | 35.42 (4.84) |  |  |  | 33.26 (5.17) |

| **HIGH**  36.25 (3.68) | | **Density *D. bunnanda*** | | | | |
| --- | --- | --- | --- | --- | --- | --- |
|  |  | **0** | **6** | **12** | **18** | **24** |
| **Density *D. birchii*** | **0** |  |  | 37.57 (3.00) |  | 35.25 (4.11) |
|  | **6** |  | 35.78 (4.06) |  | 36.32 (3.42) |  |
|  | **12** |  |  | 35.89 (3.58) |  |  |
|  | **18** |  | 36.38 (4.04) |  |  |  |
|  | **24** |  |  |  |  | 36.03 (3.18) |

(C) **(ii) Emergence time (days) females**

***Drosophila birchii***

***Drosophila bunnanda***

| **LOW**  1.63 (0.10) | | **Density *D. bunnanda*** | | | | |
| --- | --- | --- | --- | --- | --- | --- |
|  |  | **0** | **6** | **12** | **18** | **24** |
| **Density *D. birchii*** | **0** |  |  |  |  |  |
|  | **6** | 1.69 (0.09) | 1.65 (0.10) |  | 1.62 (0.10) |  |
|  | **12** | 1.63 (0.10) |  | 1.61 (0.10) |  |  |
|  | **18** |  | 1.60 (0.10) |  |  |  |
|  | **24** | 1.58 (0.10) |  |  |  | 1.55 (0.08) |

| **LOW**  1.52 (0.10) | | **Density *D. bunnanda*** | | | | |
| --- | --- | --- | --- | --- | --- | --- |
|  |  | **0** | **6** | **12** | **18** | **24** |
| **Density *D. birchii*** | **0** |  |  | 1.56 (0.09) |  | 1.52 (0.09) |
|  | **6** |  | 1.54 (0.09) |  | 1.51 (0.10) |  |
|  | **12** |  |  | 1.51 (0.10) |  |  |
|  | **18** |  | 1.52 (0.10) |  |  |  |
|  | **24** |  |  |  |  | 1.46 (0.09) |

| **MID**  1.68 (0.11) | | **Density *D. bunnanda*** | | | | |
| --- | --- | --- | --- | --- | --- | --- |
|  |  | **0** | **6** | **12** | **18** | **24** |
| **Density *D. birchii*** | **0** |  |  |  |  |  |
|  | **6** | 1.71 (0.11) | 1.71 (0.10) |  | 1.63 (0.12) |  |
|  | **12** | 1.69 (0.11) |  | 1.69 (0.12) |  |  |
|  | **18** |  | 1.64 (0.13) |  |  |  |
|  | **24** | 1.65 (0.12) |  |  |  | 1.61 (0.12) |

| **MID**  1.53 (0.11) | | **Density *D. bunnanda*** | | | | |
| --- | --- | --- | --- | --- | --- | --- |
|  |  | **0** | **6** | **12** | **18** | **24** |
| **Density *D. birchii*** | **0** |  |  | 1.55 (0.09) |  | 1.50 (0.12) |
|  | **6** |  | 1.57 (0.10) |  | 1.53 (0.11) |  |
|  | **12** |  |  | 1.51 (0.09) |  |  |
|  | **18** |  | 1.54 (0.13) |  |  |  |
|  | **24** |  |  |  |  | 1.49 (0.11) |

| **HIGH**  1.81 (0.11) | | **Density *D. bunnanda*** | | | | |
| --- | --- | --- | --- | --- | --- | --- |
|  |  | **0** | **6** | **12** | **18** | **24** |
| **Density *D. birchii*** | **0** |  |  |  |  |  |
|  | **6** | 1.85 (0.10) | 1.81 (0.11) |  | 1.82 (0.11) |  |
|  | **12** | 1.83 (0.11) |  | 1.77 (0.11) |  |  |
|  | **18** |  | 1.75 (0.11) |  |  |  |
|  | **24** | 1.78 (0.12) |  |  |  | 1.72 (0.07) |

| **HIGH**  1.67 (0.08) | | **Density *D. bunnanda*** | | | | |
| --- | --- | --- | --- | --- | --- | --- |
|  |  | **0** | **6** | **12** | **18** | **24** |
| **Density *D. birchii*** | **0** |  |  | 1.74 (0.08) |  | 1.67 (0.09) |
|  | **6** |  | 1.68 (0.07) |  | 1.66 (0.08) |  |
|  | **12** |  |  | 1.66 (0.06) |  |  |
|  | **18** |  | 1.60 (0.09) |  |  |  |
|  | **24** |  |  |  |  | 1.59 (0.06) |

(D) **(i) Wing size (mm) males**

***Drosophila birchii***

***Drosophila bunnanda***

| **LOW**  1.75 (0.12) | | **Density *D. bunnanda*** | | | | |
| --- | --- | --- | --- | --- | --- | --- |
|  |  | **0** | **6** | **12** | **18** | **24** |
| **Density *D. birchii*** | **0** |  |  |  |  |  |
|  | **6** | 1.79 (0.11) | 1.75 (0.12) |  | 1.75 (0.11) |  |
|  | **12** | 1.75 (0.11) |  | 1.76 (0.10) |  |  |
|  | **18** |  | 1.74 (0.11) |  |  |  |
|  | **24** | 1.69 (0.12) |  |  |  | 1.72 (0.12) |

| **LOW**  1.64 (0.13) | | **Density *D. bunnanda*** | | | | |
| --- | --- | --- | --- | --- | --- | --- |
|  |  | **0** | **6** | **12** | **18** | **24** |
| **Density *D. birchii*** | **0** |  |  | 1.68 (0.11) |  | 1.60 (0.14) |
|  | **6** |  | 1.68 (0.13) |  | 1.61 (0.13) |  |
|  | **12** |  |  | 1.63 (0.13) |  |  |
|  | **18** |  | 1.63 (0.13) |  |  |  |
|  | **24** |  |  |  |  | 1.53 (0.12) |

| **MID**  1.76 (0.12) | | **Density *D. bunnanda*** | | | | |
| --- | --- | --- | --- | --- | --- | --- |
|  |  | **0** | **6** | **12** | **18** | **24** |
| **Density *D. birchii*** | **0** |  |  |  |  |  |
|  | **6** | 1.81 (0.11) | 1.78 (0.11) |  | 1.72 (0.14) |  |
|  | **12** | 1.77 (0.11) |  | 1.72 (0.13) |  |  |
|  | **18** |  | 1.73 (0.13) |  |  |  |
|  | **24** | 1.73 (0.12) |  |  |  | 1.70 (0.13) |

| **MID**  1.64 (0.12) | | **Density *D. bunnanda*** | | | | |
| --- | --- | --- | --- | --- | --- | --- |
|  |  | **0** | **6** | **12** | **18** | **24** |
| **Density *D. birchii*** | **0** |  |  | 1.68 (0.11) |  | 1.57 (0.10) |
|  | **6** |  | 1.68 (0.11) |  | 1.61 (0.12) |  |
|  | **12** |  |  | 1.62 (0.12) |  |  |
|  | **18** |  | 1.66 (0.12) |  |  |  |
|  | **24** |  |  |  |  | 1.63 (0.10) |

| **HIGH**  1.92 (0.13) | | **Density *D. bunnanda*** | | | | |
| --- | --- | --- | --- | --- | --- | --- |
|  |  | **0** | **6** | **12** | **18** | **24** |
| **Density *D. birchii*** | **0** |  |  |  |  |  |
|  | **6** | 1.98 (0.10) | 1.90 (0.14) |  | 1.87 (0.15) |  |
|  | **12** | 1.93 (0.15) |  | 1.88 (0.14) |  |  |
|  | **18** |  | 1.89 (0.11) |  |  |  |
|  | **24** | 1.87 (0.11) |  |  |  | 1.80 (0.10) |

| **HIGH**  1.78 (0.11) | | **Density *D. bunnanda*** | | | | |
| --- | --- | --- | --- | --- | --- | --- |
|  |  | **0** | **6** | **12** | **18** | **24** |
| **Density *D. birchii*** | **0** |  |  | 1.85 (0.07) |  | 1.74 (0.11) |
|  | **6** |  | 1.79 (0.09) |  | 1.76 (0.13) |  |
|  | **12** |  |  | 1.74 (0.10) |  |  |
|  | **18** |  | 1.74 (0.13) |  |  |  |
|  | **24** |  |  |  |  | 1.74 (0.10) |

(D) **(ii) Wing size (mm) females**

**Table S3.** Variance components for full models (from Table 1)

| (A) **Productivity** | | | | | |
| --- | --- | --- | --- | --- | --- |
| **Species** |  | **Variance component** | **Variance** | **χ^2^_(1 df)_** | ***P*** |
| ***Drosophila birchii*** | | Source population | 0.000 | 0 | 1 |
|  |  | Residual | 0.304 |  |  |
| ***Drosophila bunnanda*** | | *Source population* | *9.63 x 10^-3^* | *5.328* | *0.021* |
|  |  | Residual | 0.178 |  |  |
| (B) **Sex ratio** | | | | | |
| **Species** |  | **Variance component** | **Variance** | **χ^2^_(1 df)_** | ***P*** |
| ***Drosophila birchii*** | | Source population | 0.001 | 0.448 | 0.503 |
|  |  | Residual | 0.020 |  |  |
| ***Drosophila bunnanda*** | | Source population | 1.53 x 10^-2^ | 0 | 1 |
|  |  | Residual | 5.23 x 10^-9^ |  |  |
| (C) **Emergence time** | | | | | |
| **Species** | **Sex** | **Variance component** | **Variance** | **χ^2^_(1 df)_** | ***P*** |
| ***Drosophila birchii*** | **Male** | Source population | 0.006 | 0 | 1 |
|  |  | *Vial* | *5.365* | *400.22* | *<2.2 x 10^-16^* |
|  |  | Residual | 31.95 |  |  |
|  | **Female** | Source population | 0.106 | 1.293 | 0.256 |
|  |  | *Vial* | *4.818* | *368.22* | *<2.2 x 10^-16^* |
|  |  | Residual | 31.91 |  |  |
| ***Drosophila bunnanda*** | **Male** | Source population | 0.000 | 0 | 1 |
|  |  | *Vial* | *3.473* | *90.88* | *<2.2 x 10^-16^* |
|  |  | Residual | 25.62 |  |  |
|  | **Female** | *Source population* | *0.314* | *4.481* | *0.034* |
|  |  | *Vial* | *3.294* | *113.86* | *<2.2 x 10^-16^* |
|  |  | Residual | 26.88 |  |  |
| (D) **Wing size** | | | | | |
| **Species** | **Sex** | **Variance component** | **Variance** | **χ^2^_(1 df)_** | ***P*** |
| ***Drosophila birchii*** | **Male** | *Source population* | *1.31 x 10^-4^* | *9.223* | *0.002* |
|  |  | *Vial* | *2.29 x 10^-3^* | *519.58* | *<2.2 x 10^-16^* |
|  |  | Residual | 8.26 x 10^-3^ |  |  |
|  | **Female** | *Source population* | *2.51 x 10^-4^* | *11.38* | *7.43 x 10^-4^* |
|  |  | *Vial* | 3.30 x 10^-3^ | 326.17 | *<2.2 x 10^-16^* |
|  |  | Residual | 9.88 x 10^-3^ |  |  |
| ***Drosophila bunnanda*** | **Male** | *Source population* | *1.43 x 10^--4^* | *4.840* | *0.028* |
|  |  | *Vial* | *1.49 x 10^-3^* | *81.66* | *<2.2 x 10^-16^* |
|  |  | Residual | 7.86 x 10^-3^ |  |  |
|  | **Female** | *Source population* | *3.45 x 10^-4^* | *8.656* | *0.003* |
|  |  | *Vial* | *2.06 x 10^-3^* | *44.20* | *2.97 x 10^-11^* |
|  |  | Residual | 1.21 x 10^-2^ |  |  |

**Table S4.** Variance components for models run separately for each elevation (from Table S1)

| (A) **Productivity** | | | | | | | |
| --- | --- | --- | --- | --- | --- | --- | --- |
| **Species** | | **Elevation** |  | **Variance component** | **Variance** | **χ^2^_(1 df)_** | ***P*** |
| ***Drosophila birchii*** | | **Low** |  | Source population | 0.000 | 2.815 | 0.093 |
|  |  |  |  | Residual | 0.389 |  |  |
|  |  | **Mid** |  | Source population | 0.000 | 0 | 1 |
|  |  |  |  | Residual | 0.250 |  |  |
|  |  | **High** |  | Source population | 0.003 | 0 | 1 |
|  |  |  |  | Residual | 0.272 |  |  |
| ***Drosophila bunnanda*** | | **Low** |  | Source population | 0.016 | 0.410 | 0.522 |
|  |  |  |  | Residual | 0.245 |  |  |
|  |  | **Mid** |  | Source population | 0.006 | 0 | 1 |
|  |  |  |  | Residual | 0.150 |  |  |
|  |  | **High** |  | *Source population* | *0.017* | *6.559* | *0.010* |
|  |  |  |  | Residual | 0.130 |  |  |
| (B) **Sex ratio** | | | | | | | |
| **Species** | | **Elevation** |  | **Variance component** | **Variance** | **χ^2^_(1 df)_** | ***P*** |
| ***Drosophila birchii*** | | **Low** |  | Source population | 1.30 x 10^-9^ | 0 | 1 |
|  |  |  |  | Residual | 0.024 |  |  |
|  |  | **Mid** |  | Source population | 0.008 | 1.421 | 0.233 |
|  |  |  |  | Residual | 0.010 |  |  |
|  |  | **High** |  | Source population | 0.000 | 0 | 1 |
|  |  |  |  | Residual | 5.92 x 10^-17^ |  |  |
| ***Drosophila bunnanda*** | | **Low** |  | Source population | 0.015 | 0 | 1 |
|  |  |  |  | Residual | 1.40 x 10^-15^ |  |  |
|  |  | **Mid** |  | Source population | 8.55 x 10^-9^ | 0 | 1 |
|  |  |  |  | Residual | 4.60 x 10^-2^ |  |  |
|  |  | **High** |  | Source population | 0.034 | 1.82 | 0.177 |
|  |  |  |  | Residual | 2.60 x 10^-9^ |  |  |
| (C) **Emergence time** | | | | | | | |
| **Species** | | **Elevation** | **Sex** | **Variance component** | **Variance** | **χ^2^_(1 df)_** | ***P*** |
| ***Drosophila birchii*** | | **Low** | **Male** | Source population | 0.169 | 0.472 | 0.492 |
|  |  |  |  | *Vial* | *6.261* | *299.74* | *<2.2 x 10^-16^* |
|  |  |  |  | Residual | 33.11 |  |  |
|  |  |  | **Female** | Source population | 0.210 | 0.996 | 0.318 |
|  |  |  |  | *Vial* | *5.438* | *237.50* | *<2.2 x 10^-16^* |
|  |  |  |  | Residual | 34.23 |  |  |
|  |  | **Mid** | **Male** | Source population | 0.000 | 0 | 1 |
|  |  |  |  | *Vial* | *4.261* | *51.11* | *8.75 x 10^-13^* |
|  |  |  |  | Residual | 35.64 |  |  |
|  |  |  | **Female** | Source population | 0.112 | 0.081 | 0.777 |
|  |  |  |  | *Vial* | *4.852* | *81.97* | *<2.2 x 10^-16^* |
|  |  |  |  | Residual | 32.34 |  |  |
|  |  | **High** | **Male** | Source population | 0.000 | 0 | 1 |
|  |  |  |  | *Vial* | *5.360* | *54.78* | *1.35 x 10^-13^* |
|  |  |  |  | Residual | 16.88 |  |  |
|  |  |  | **Female** | Source population | 0.000 | 0 | 1 |
|  |  |  |  | *Vial* | *4.062* | *47.17* | *6.52 x 10^-12^* |
|  |  |  |  | Residual | 20.07 |  |  |
| ***Drosophila bunnanda*** | | **Low** | **Male** | Source population | 0.000 | 0 | 1 |
|  |  |  |  | *Vial* | *5.253* | *66.40* | *3.68 x 10^-16^* |
|  |  |  |  | Residual | 31.80 |  |  |
|  |  |  | **Female** | Source population | 1.93 x 10^-12^ | 0 | 1 |
|  |  |  |  | *Vial* | *4.421* | *74.32* | *<2.2 x 10^-16^* |
|  |  |  |  | Residual | 33.06 |  |  |
|  |  | **Mid** | **Male** | Source population | 0.036 | 0 | 1 |
|  |  |  |  | *Vial* | *2.372* | *11.22* | *8.10 x 10^-4^* |
|  |  |  |  | Residual | 20.91 |  |  |
|  |  |  | **Female** | *Source population* | *1.172* | *7.280* | *6.97 x 10^-3^* |
|  |  |  |  | *Vial* | *2.845* | *16.89* | *3.96 x 10^-5^* |
|  |  |  |  | Residual | 22.41 |  |  |
|  |  | **High** | **Male** | Source population | 0.000 | 0 | 1 |
|  |  |  |  | *Vial* | *2.140* | *13.30* | *2.66 x 10^-4^* |
|  |  |  |  | Residual | 11.81 |  |  |
|  |  |  | **Female** | Source population | 0.000 | 0 | 1 |
|  |  |  |  | *Vial* | *2.561* | *44.31* | *2.80 x 10^-11^* |
|  |  |  |  | Residual | 10.61 |  |  |
| (D) **Wing size** | | | | | | | |
| **Species** | | **Elevation** | **Sex** | **Variance component** | **Variance** | **χ^2^_(1 df)_** | ***P*** |
| ***Drosophila birchii*** | | **Low** | **Male** | *Source population* | *1.77 x 10^-4^* | *5.725* | *0.017* |
|  |  |  |  | *Vial* | *1.81 x 10^-3^* | *358.99* | *<2.2 x 10^-16^* |
|  |  |  |  | Residual | 7.49 x 10^-3^ |  |  |
|  |  |  | **Female** | *Source population* | *3.29 x 10^-4^* | *5.182* | *0.023* |
|  |  |  |  | *Vial* | *2.63 x 10^-3^* | *163.43* | *<2.2 x 10^-16^* |
|  |  |  |  | Residual | 9.72 x 10^-3^ |  |  |
|  |  | **Mid** | **Male** | Source population | 1.67 x 10^-17^ | 0 | 1 |
|  |  |  |  | *Vial* | *2.33 x 10^-3^* | *92.61* | *<2.2 x 10^-16^* |
|  |  |  |  | Residual | 1.02 x 10^-2^ |  |  |
|  |  |  | **Female** | Source population | 2.90 x 10^-5^ | 0 | 1 |
|  |  |  |  | *Vial* | *3.70 x 10^-3^* | *104.79* | *<2.2 x 10^-16^* |
|  |  |  |  | Residual | 9.77 x 10^-3^ |  |  |
|  |  | **High** | **Male** | Source population | 2.97 x 10^-4^ | 1.853 | 0.173 |
|  |  |  |  | *Vial* | *3.65 x 10^-3^* | *63.17* | *1.89 x 10^-15^* |
|  |  |  |  | Residual | 7.98 x 10^-3^ |  |  |
|  |  |  | **Female** | Source population | 5.95 x 10^-4^ | 2.981 | 0.084 |
|  |  |  |  | *Vial* | *4.07 x 10^-3^* | *40.10* | *2.41 x 10^-10^* |
|  |  |  |  | Residual | 1.07 x 10^-2^ |  |  |
| ***Drosophila bunnanda*** | | **Low** | **Male** | *Source population* | *4.33 x 10^-4^* | *9.493* | *2.06 x 10^-3^* |
|  |  |  |  | *Vial* | *1.22 x 10^-3^* | *40.76* | *1.72 x 10^-10^* |
|  |  |  |  | Residual | 8.03 x 10^-3^ |  |  |
|  |  |  | **Female** | *Source population* | *5.79 x 10^-4^* | *5.236* | *0.022* |
|  |  |  |  | *Vial* | *2.31 x 10^-3^* | *27.56* | *1.52 x 10^-7^* |
|  |  |  |  | Residual | 1.35 x 10^-2^ |  |  |
|  |  | **Mid** | **Male** | Source population | 0.000 | 0 | 1 |
|  |  |  |  | *Vial* | *2.34 x 10^-3^* | *22.09* | *2.60 x 10^-6^* |
|  |  |  |  | Residual | 9.23 x 10^-3^ |  |  |
|  |  |  | **Female** | Source population | 0.000 | 0 | 1 |
|  |  |  |  | Vial | 1.28 x 10^-3^ | 3.368 | 0.066 |
|  |  |  |  | Residual | 1.18 x 10^-2^ |  |  |
|  |  | **High** | **Male** | Source population | 1.48 x 10^-4^ | 0.848 | 0.357 |
|  |  |  |  | *Vial* | *8.59 x 10^-4^* | *7.508* | *0.006* |
|  |  |  |  | Residual | 4.83 x 10^-3^ |  |  |
|  |  |  | **Female** | Source population | 6.52 x 10^-4^ | 2.655 | 0.103 |
|  |  |  |  | *Vial* | *2.58 x 10^-3^* | *13.90* | *1.93 x 10^-4^* |
|  |  |  |  | Residual | 8.15 x 10^-3^ |  |  |
